# Arabidopsis Casein Kinase 2 triggers Stem Cell Exhaustion under Al Toxicity and Phosphate Deficiency Through activation of the DNA Damage Response pathway

**DOI:** 10.1101/2020.11.30.404459

**Authors:** Pengliang Wei, Manon Demulder, Pascale David, Thomas Eekhout, Kaoru Okamoto Yoshiyama, Long Nguyen, Ilse Vercauteren, Dominique Eeckhout, Margot Galle, Geert De Jaeger, Paul Larsen, Dominique Audenaert, Thierry Desnos, Laurent Nussaume, Remy Loris, Lieven De Veylder

## Abstract

Aluminum (Al) toxicity and inorganic phosphate (Pi) limitation are widespread chronic abiotic and mutually enhancing stresses that profoundly affect crop yield. Both stresses cause a strong inhibition of root growth, resulting from a progressive exhaustion of the stem cell niche. Here, we report on a casein kinase 2 (CK2) inhibitor identified by its capability to maintain a functional root stem cell niche under Al toxic conditions. CK2 operates through phosphorylation of the cell cycle checkpoint activator SUPPRESSOR OF GAMMA RADIATION1 (SOG1), priming its activity under DNA-damaging conditions. In addition to yielding Al tolerance, CK2 and SOG1 inactivation prevents meristem exhaustion under Pi starvation, revealing the existence of a low Pi-induced cell cycle checkpoint that depends on the DNA damage activator ATAXIA-TELANGIECTASIA MUTATED. Overall, our data reveal an important physiological role for the plant DNA damage response pathway under agriculturally limiting growth conditions, opening new avenues to cope with Pi limitation.

**ONE-SENTENCE SUMMARY:** Casein kinase 2 and DNA damage response regulators play a pivotal role in the control of Arabidopsis root growth in response to Al toxicity and phosphate limitation.

## INTRODUCTION

As a major constituent of soils worldwide, aluminum (Al) is considered to be an important limitation to agricultural productivity (Von Uexküll and Mutert, 1995; Bot et al., 2000). Al is normally locked in an insoluble complex within minerals and does not cause toxicity to plants. However, in acid soil environments, which occur both naturally and as a result of pollution and land management practices (Vitorello et al., 2005; Kochian et al., 2015), Al becomes increasingly soluble and speciates to the toxic Al^3+^ form. The most evident consequence of Al toxicity is root growth inhibition, which occurs even at low micromolar concentrations. As a result of this inhibition, nutrient uptake is reduced, resulting in nutritional deficiency in shoots and leaves that cause an overall reduction in plant growth and yield.

Internalized Al^3+^ interacts promiscuously with both inter- and intracellular anionic sites, including cell wall components, the plasma membrane and mitochondria (Kopittke et al., 2016). Much of the internalized Al^3+^ is found in the cell wall bound to pectin, where it affects cell wall rigidity and reduces cell elongation in conjunction with causing gross physical damage (Ma et al., 2004; Horst et al., 2010). Al^3+^ also accumulates within the cell and, unless properly sequestered in the vacuole or transported to less-sensitive tissues, can cause severe consequences that lead to termination of cell division and root growth. A major intracellular target of Al^3+^ is the DNA, where noncovalent binding of Al^3+^ to the negatively charged phosphodiester backbone is thought to result in a conformational change from the B-form to Z-DNA, resulting in increased DNA rigidity that leads to difficulty in unwinding during DNA replication (Latha et al., 2002). In addition, Al^3+^ inhibits DNA synthesis and causes DNA fragmentation and the generation of micronuclei (Kopittke et al., 2016). As a result, the cells within the root meristem lose their stem cells and division competence (Sjogren et al., 2015), a process known as meristem exhaustion. Strikingly, this exhaustion can be reversed by mutations in DNA damage response (DDR) regulators, including mutations within the *SUPPRESSOR OF GAMMA RESPONSE 1 (SOG1)* gene (Sjogren et al., 2015) that has been put forward as the plant counterpart of the mammalian p53 tumor suppressor protein (Yoshiyama et al., 2009; Yoshiyama et al., 2014). *SOG1* encodes a transcription factor of the NAC domain family and is responsible for the transcriptional induction of DNA damage-responsive genes operating downstream of the ATAXIA-TELANGIECTASIA MUTATED (ATM) and ATM-RELATED (ATR) kinases (Hu et al., 2016; Bourbousse et al., 2018; Ogita et al., 2018). Activation of SOG1 in response to DNA double-strand breaks (DSBs) requires its phosphorylation by ATM on consensus SQ amino-acid motifs (Yoshiyama et al., 2013). In addition. SOG1 is likely also an *in vivo* phosphorylation target of ATR (Sjogren et al., 2015).

Al toxicity is intrinsically linked to the agricultural problem of phosphorus (P)-limitation, which can be explained by the immobilization of P through its chelation with Al^3+^, reducing the bioavailability of inorganic phosphate (Pi) to plants (Raghothama, 1999; Chiou and Lin, 2011). Strikingly, the symptoms of Pi deficiency resemble those of Al toxicity, including overall root stunting, cell wall stiffening, callose accumulation and meristem exhaustion (Müller et al., 2015; Balzergue et al., 2017). Correspondingly, some of the Arabidopsis mutants that are hypersensitive to Al show a similar strong growth defect under P-limiting conditions. These include plants being deficient for ALUMINUM SENSITIVE1 (ALS1) or ALUMINUM-SENSITIVE3 (ALS3), both encoding ABC transporters with a role in the sequestration of Al into the vacuole, as well as mutants for the SENSITIVITY TO PROTON RHIZOTOXICITY1 (STOP1) transcription factor and its target *ALUMINUM-ACTIVATED MALATE TRANSPORTER1* (*ALMT1*), which encodes a malate efflux channel (Balzergue et al., 2017; Mora-Macias et al., 2017; Godon et al., 2019). Interestingly, the root growth phenotypes observed under Pi deficiency appear to be iron (Fe) dependent. Under low P conditions, labile Fe^3+^ ions accumulate in the root tip, probably causing buildup of reactive oxygen species (ROS) and peroxidase activity that stiffen the cell wall and cause callose accumulation, interfering with the movement of stem cell regulators between neighboring cells (Svistoonoff et al., 2007; Müller et al., 2015; Balzergue et al., 2017; Wang et al., 2019).

At the biochemical level, plants respond to P starvation through activation of PHOSPHATE STARVATION RESPONSE 1 (PHR1) and related MYB transcription factors, inducing Pi scavenging and transport activities, membrane lipid remodeling, increase of root-to-shoot growth ratio, and suppression of photosynthesis and photorespiration. Pi uptake is promoted through accumulation of high-affinity Pi transporter PHT1, as well as of PHOSPHATE TRANSPORTER TRAFFIC FACILITATOR 1 (PHF1) (González et al., 2005; Bayle et al., 2011; Nussaume et al., 2011), an ER exit cofactor that enables PHT1 traffic to the plasma membrane. Previously, casein kinase 2 (CK2) has been identified as a new factor controlling PHT1 availability at the plasma membrane (Chen et al., 2015). CK2 represents an evolutionary conserved S/T kinase involved in a wide variety of cellular functions (Meggio and Pinna, 2003; Mulekar and Huq, 2014). The CK2 holoenzyme consists of a tetramer consisting of two α subunits and two regulatory β subunits, with each type of subunit encoded by four different genes in *Arabidopsis*. The CK2α subunits are ubiquitously expressed throughout development but show variability in their subcellular localization patterns, with CKA1 (*At5g67380*), CKA2 (*At3g50000*) and CKA3 (*At2g23080*) being predominantly localized in the nucleus, whereas CKA4 (*At2g23070*) localizes to the chloroplasts (Salinas et al., 2006). In rice, CK2 phosphorylates PHT1 and prevents its interaction with PHF1, resulting in ER retention of PHT1. Low Pi triggers degradation of the CK2β subunits, probably resulting in a decrease in CK2 activity and thus relieving PHT1 inhibition (Chen et al., 2015).

In addition to signaling in response to Pi starvation, plant CK2 activity has also been linked to maintenance of genome integrity. Plants expressing a dominant-negative *CK2* allele were found to be hypersensitive to genotoxic stress conditions, including high radiation, UV-C and MMS treatment. Although the molecular basis of this hypersensitivity is unknown, transcriptome analysis suggested that it might originate from lower expression of DNA repair genes (Moreno-Romero et al., 2012). Likewise, blocking CK2 activity in S-phase cells was found to trigger premature DNA condensation, implying the lack of a G2/M checkpoint (Espunya et al., 1999). Combined, these data suggest a role for CK2 activity in both aspects of the DDR pathway, being the transcriptional induction of DNA damage repair genes and coordination of DNA repair with cell cycle progression through activation of a cell cycle inhibitory mechanism.

Here, we report on the identification of a chemical inhibitor of the plant CK2 kinase, isolated through a small molecule screen aimed at identifying compounds that result in Al toxicity tolerance. We demonstrate that CK2 controls the DDR pathway through phosphorylation of SOG1. Under CK2-inhibiting conditions, SOG1 cannot be phosphorylated by ATM, suggesting a role for CK2 in priming SOG1 for its activation by the ATM kinase under DNA-damaging conditions. Strikingly, we demonstrate that next to controlling Al toxicity-induced stem cell exhaustion, CK2, SOG1 and ATM control root growth in response to Pi deficiency, demonstrating that cell cycle checkpoint regulators are causative agents that induce a root growth arrest under P-limiting growth conditions.

## RESULTS

### C43 Promotes Al Resistance Without Affecting Al Uptake

The root stem cell niche holds an organizing quiescent center (QC) that is lost under Al toxic conditions (Sjogren et al., 2015). Based on this observation, a small molecule compound screen was designed (see Methods) to identify compounds that help maintain expression of the QC25 stem cell marker in the presence of Al under acidic conditions (**Figure 1**). Seeds of plants that express *β-glucuronidase* (*GUS*) under the control of the *QC25* promoter were sown in 96-well plates. After three days Al was added under acidic conditions (pH 4.2) at a final concentration of 0.5 mM in combination with chemicals (at 50 μM) from the Pharmacological Diversity Set (Enamine). Five days after treatment, seedlings were analyzed for a GUS-positive signal in the root stem cell region. In total, 10,000 different organic molecules were screened, resulting in the identification of 355 primary hit compounds. Among these, 44 compounds were confirmed in a secondary screen. Here, we report on compound C43 (N-methyl-2-(5,6,7,8-tetrahydro-[1]benzothiolo[2,3-d]pyrimidin-4-ylsulfanyl)acetamide).

**Figure 1.**
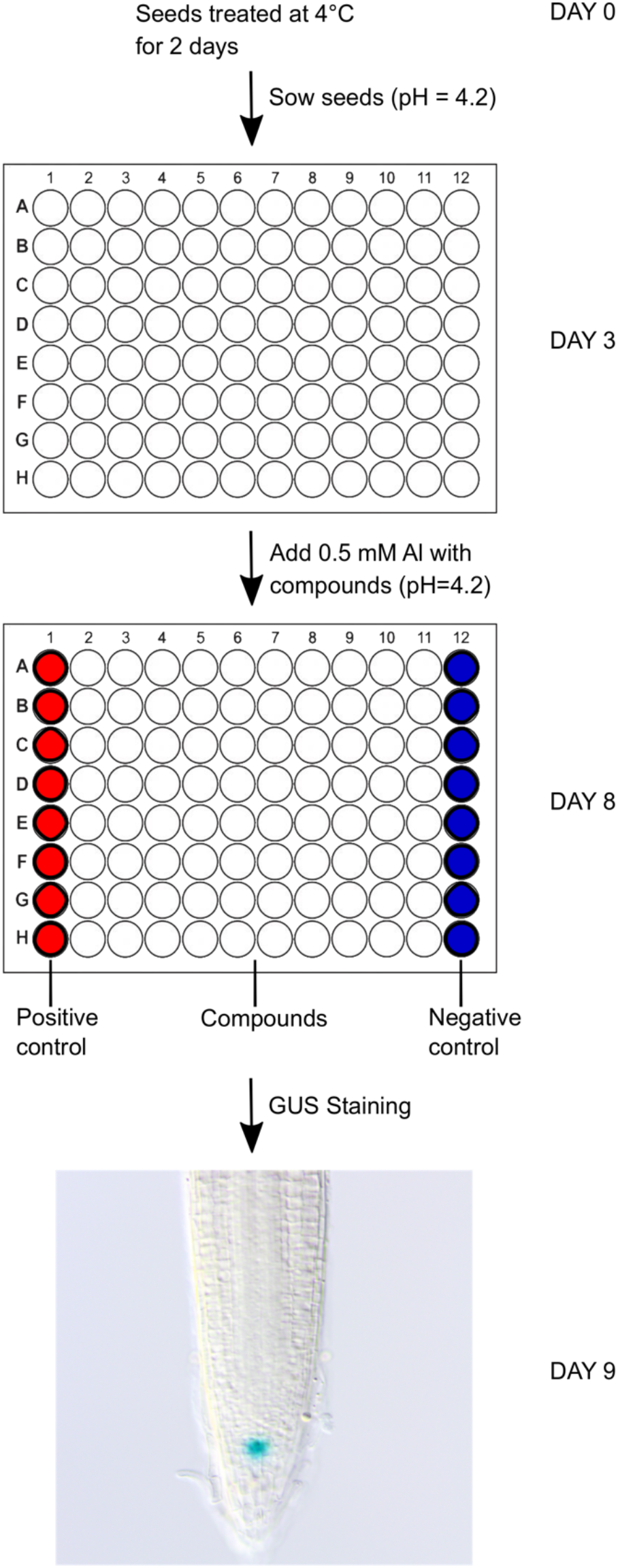
Schematic Set Up to Screen for Compounds Granting Al Toxicity Resistance

Correlated with maintenance of expression of the QC25 marker (**Figure 2A**), C43 application slightly but significantly rescued the Al-dependent growth inhibition phenotype, despite causing an inhibitory effect on roots grown in the absence of Al (**Figure 2B**). A doseresponse analysis demonstrated a maximum root growth rescue versus inhibitory activity at a concentration of 50 μM C43 (**Supplemental Figure 1**).

**Figure 2.**
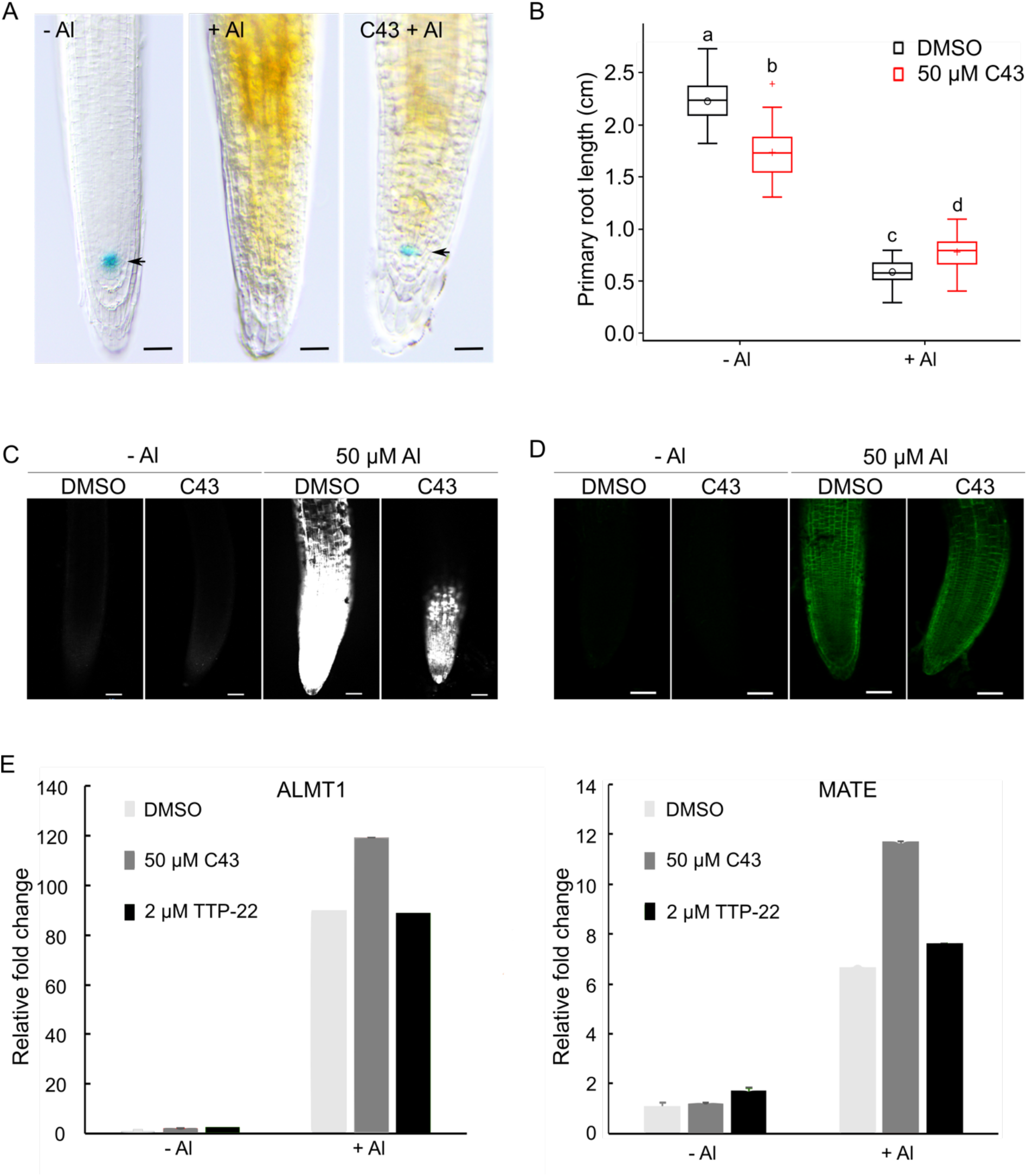
C43 Promotes Al Resistance Without Affecting Al Uptake. (A) Three-day-old *QC25-GUS* seedings grown for 5 days in the presence of 0.5 mM AlCl_3_ without (+ Al) or with 50 μM C43 (C43 +Al) for 5 days. Seedling treated without AlCl_3_ (-Al) were used as positive control. GUS staining was performed overnight. Arrows point to the position of QC cells. Bars = 50 μm. (B) Primary root length of 7-day-old wild-type (Col-0) seedlings grown in a soaked gel environment (pH 4.2) in the absence (-Al) or presence (+ Al) of 1.5 mM AlCl_3_, together with solvent control (DMSO) or 50 μM C43. Data are represented by box and whisker plots. Significant differences (p < 0.01) as determined by mixed model analysis with Tukey’s post-hoc correction are indicated by letters above bars (n = 3, with > 8 plants in each repeat). (C) Five-day-old wild-type seedlings hydroponically grown (at pH 4.2) in the presence of solvent control (DMSO) or 50 μM C43 and exposed to either 0 (control) or 50 μM AlCl_3_ for 24 h. Seedlings were stained with aniline blue and visualized using fluorescence microscopy for callose deposition. Bars = 50 μm. (D) Five-day-old wild-type seedlings hydroponically grown (at pH 4.2) in the presence of solvent control (DMSO) or 50 μM C43 and exposed to either 0 (control) or 50 μM AlCl3 for 1 h. Seedlings were Morin stained and visualized using fluorescence microscopy for Al accumulation. Bars = 50 μm. (E) Expression of Al-responsive genes *ALMT1* (left) and *MATE* (right) as measured by RT-qPCR in wild-type seedlings grown for 5 d (at pH 4.2) under control conditions (-Al) or in the presence of 1.50 mM AlCl_3_ (+ Al) together with solvent control (DMSO), 50 μM C43 or 2 μM TTP-22. The expression level of DMSO control grown in the absence of Al was arbitrary set to one. Data represent least square means ± se (n = 3, with > 50 plants in each repeat).

Callose accumulation within the root tip represents a well-known Al-induced response (Horst et al., 2010). Such accumulation was strongly reduced upon the co-application of Al with C43 (**Figure 2C**). Visualization of Al through Morin staining indicated that C43 application did not prevent uptake of Al (**Figure 2D**). Correspondingly, plants displayed an strong transcriptional activation of the Al-responsive *ALMT1* (*At1g08430*) and *MATE* (*At1g51340*) genes when grown under Al toxic conditions with and without C43 (**Figure 2E**).

### C43 Represents a Potential CK2 Inhibitor

C43 is structurally similar to compound TTP-22 (**Figure 3A**), a potent ATP-competitive inhibitor of human casein kinase 2 (CK2) (Golub et al., 2011). Identical to C43, expression of the QC25 marker was maintained following TTP-22 application in the presence of Al (**Figure 3B**), and partially rescued the Al toxicity-induced root growth inhibition (**Figure 3C**) and callose accumulation phenotypes (**Supplemental Figure 2A**) without affecting Al accumulation (**Supplemental Figure 3B**) or Al-induced *ALMT1* and *MATE* expression (**Figure 2E**). A dose-response analysis demonstrated a maximum root growth rescue versus inhibitory activity at a concentration of 2 μM TTP-22 (**Supplemental Figure 1**). Fitting the hypothesis of CK2 activity to be important in the Al toxicity response, *CK2* knockdown mutants (*cka123*; (Lu et al., 2011) displayed a partial rescue of root growth under Al toxic conditions (**Figure 3D**). Additionally, whereas both C43 or TTP-22 application rescued growth of Al-treated Col-0 plants, for the *cka123* line no significant additional increase in root growth could be observed when the chemicals were added (**Figure 3D**), suggesting that the C43- or TTP-22-dependent rescue of wild-type plants is predominantly dependent on the inhibition of CK2 activity.

**Figure 3.**
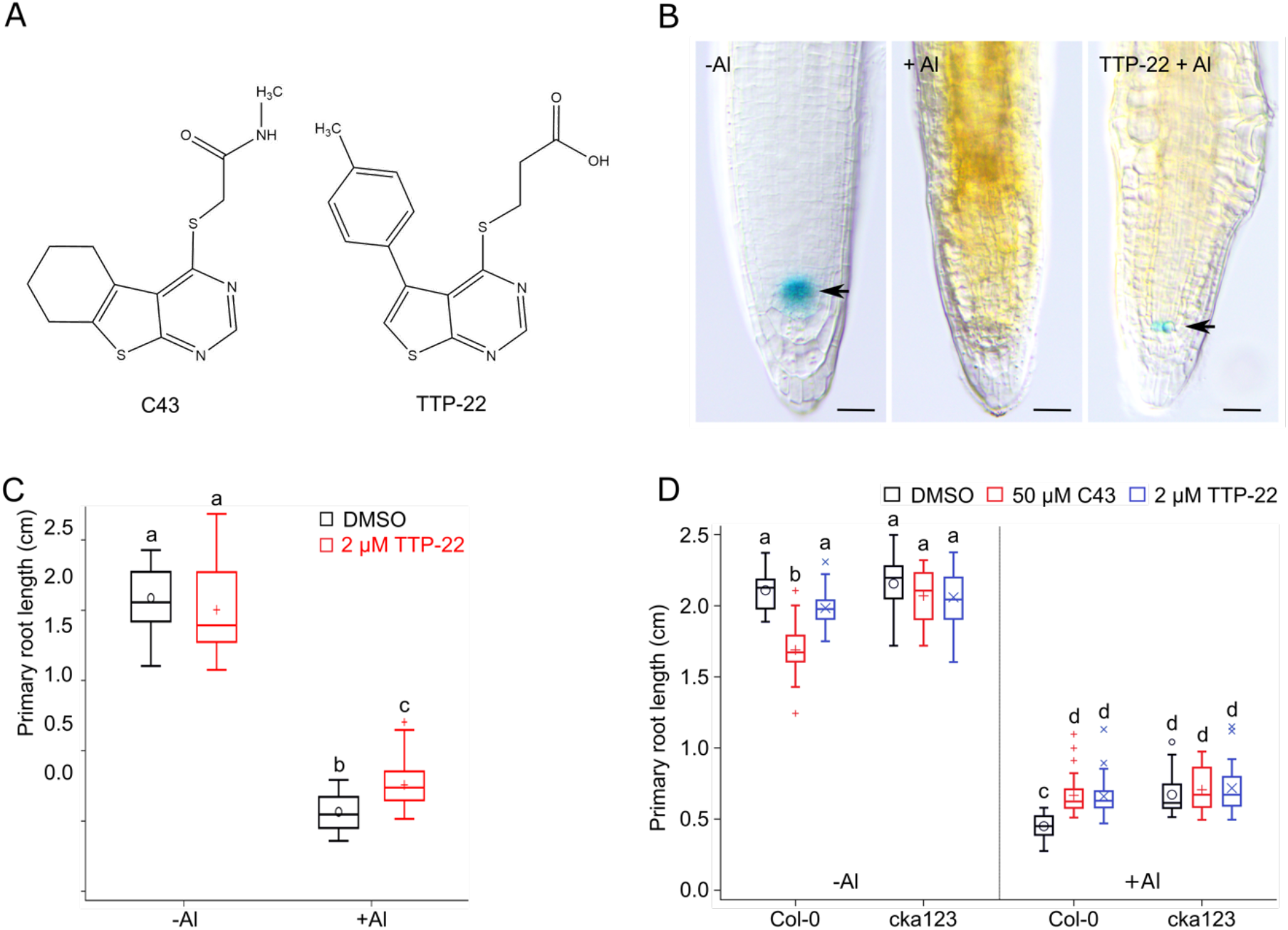
C43 Represents a Potential CK2 Inhibitor. (A) C43 and TTP-22 chemical structures. (B) Three-day-old *QC25-GUS* seedlings grown for 5 days in the presence of 0.5 mM AlCl_3_ without (+ Al) or with 2 μM TTP-22 (TTP-22 + Al). Seedling treated without AlCl_3_ (-Al) were used as positive control. GUS staining was performed overnight. Arrows point to the position of QC cells. Bars = 50 μm. (C) Primary root length of 7-day-old wild-type (Col-0) plants grown in a soaked gel environment (pH 4.2) in the absence (-Al) or presence (+ Al) of 1.5 mM AlCl_3_, together with solvent control (DMSO) or 2 μM TTP-22. Data are represented by box and whisker plots. Significant differences (p < 0.01) as determined by mixed model analysis with Tukey’s post-hoc correction are indicated by letters above bars (n = 3, with > 8 plants in each repeat). (D) Primary root length of 7-day-old wild-type and *cka123* plants grown in a soaked gel environment (pH 4.2) in the absence (-Al) or presence (+ Al) of 1.5 mM AlCl_3_, together with solvent control (DMSO), 50 μM C43 or 2 μM TTP-22. Data are represented by box and whisker plots. Significant differences (p < 0.01) as determined by mixed model analysis with Tukey’s post-hoc correction are indicated by letters above bars (n = 3, with > 8 plants in each repeat).

As support for the identified drugs to act as a plant CK2 inhibitor, binding to CK2 was tested through a thermal shift assay using purified Arabidopsis CKA1 protein. Upon addition of TTP-22, a stabilization of CKA1 towards thermal unfolding was observed in a concentration-dependent manner (**Supplemental Figure 3A**). Likewise, adding TTP-22 reduced CKA1 activity towards the substrate HD2B (**Supplemental Figure 3B**). Differently from TTP-22, we failed to detect CKA1 inhibition or binding for C43, probably due to its lower solubility and activity.

In order to confirm that TTP-22 physically interacts with Arabidopsis CKA1, co-crystallization assays were carried out. Crystals could be found to diffract to 1.8Å with a clear extra electron density pattern in the active site cleft that corresponds to TTP-22 (**Figures 4A and 4B**). The heterocyclic ring of TTP-22 is enclosed between two hydrophobic patches composed of hydrophobic side chains of I61 and V48 on one side with M158 and I169 on the other (**Figure 4C**). The CKA1-TTP-22 interaction depends on Van der Waals interactions with the hydrophobic patches formed by the two beta sheets composed of respectively strands β1-2 and β7-9, Hydrogen bonds occur between the nitrogen of the TTP-22 heterocyclic ring with the backbone NH of V111 (**Figure 4C**). Other crucial hydrogen bonds are formed between TTP-22 and the backbone nitrogen of K63 and the side-chain NH_2_ group of D170. TTP-22 is vested in the catalytic cleft through multiple interactions with surrounding residues and is thus likely to block CK2 activity by precluding ATP binding.

**Figure 4.**
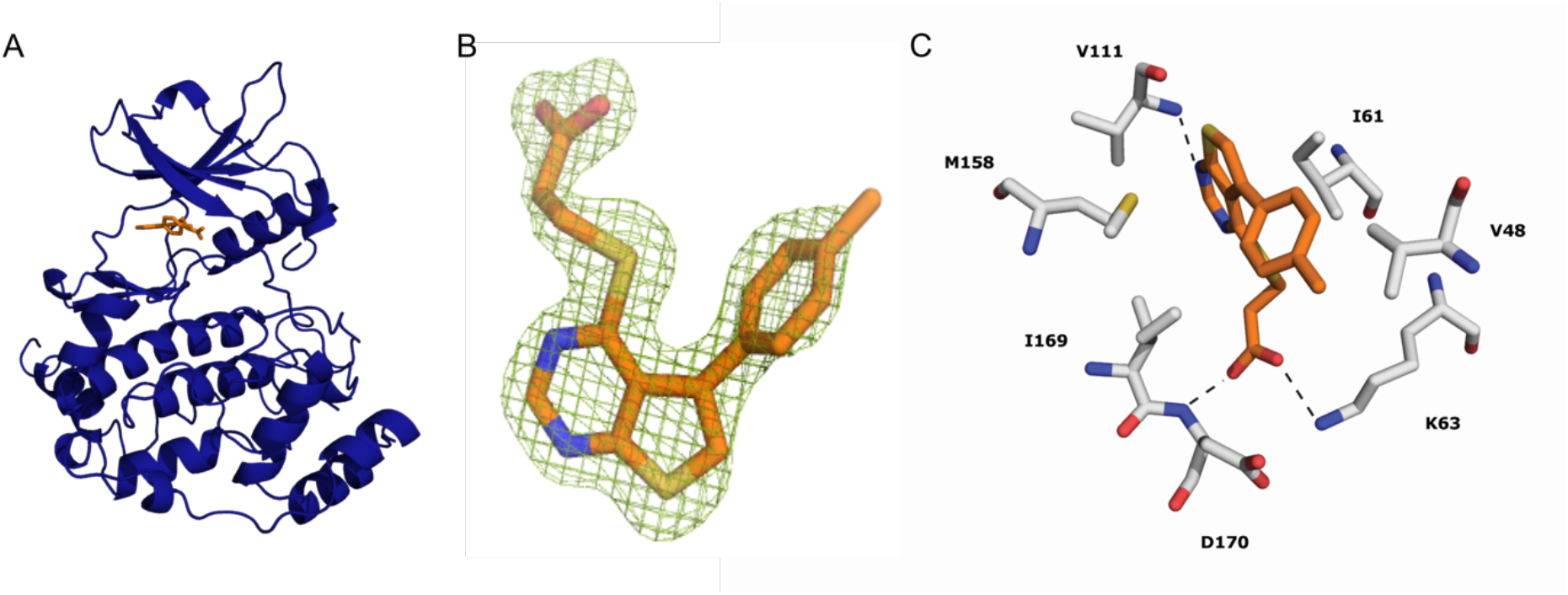
TTP-22 Binds the Catalytic Cleft of *A. thaliana* CKA1. (A) Cartoon representation of CKA1 in complex with TTP-22 (orange). (B) OMIT electron density of TTP-22 shown at 4σ. (C) Close-up of the catalytic cleft showing the interactions of TTP-22 with the protein. Residues are labeled and colored according to atom type. Carbon atoms are grey for protein and orange for TTP-22, whereas oxygen atoms are shown in red, nitrogen atoms in blue and sulfur atoms in yellow. Hydrogen bonds between TTP-22 and the active site residues are shown with black dashed lines.

### CK2 Operates Upstream of SOG1

Previously, knockdown of Arabidopsis CK2 activity has been correlated with increased DNA damage sensitivity (Moreno-Romero et al., 2012), while loss-of-function mutants for the Arabidopsis SOG1 DNA damage checkpoint regulator show Al tolerance (Sjogren et al., 2015). Therefore, it was tested whether CK2 and SOG1 operate in the same pathway. As reported, SOG1-deficient plants displayed a better growth performance compared to control plants under Al toxic conditions (**Figure 5A**). However, in contrast to wild-type plants, no additional increase in root growth could be observed when C43 was added to *sog1-7* mutant plants in the presence of Al (**Figure 5A**). These data suggest that CK2 operates in a SOG1-dependent manner. In support of this, the *cka123* line displayed increased sensitivity to the replication fork inhibitor hydroxyurea (HU) (**Figure 5B**) similar to what was reported for *sog1* mutants (Hu et al., 2015). Also application of C43 or TTP-22 resulted in increased HU sensitivity (**Figure 5C**) for wild-type Arabidopsis, with this being dependent on SOG1 activity (**Figure 5D**).

**Figure 5.**
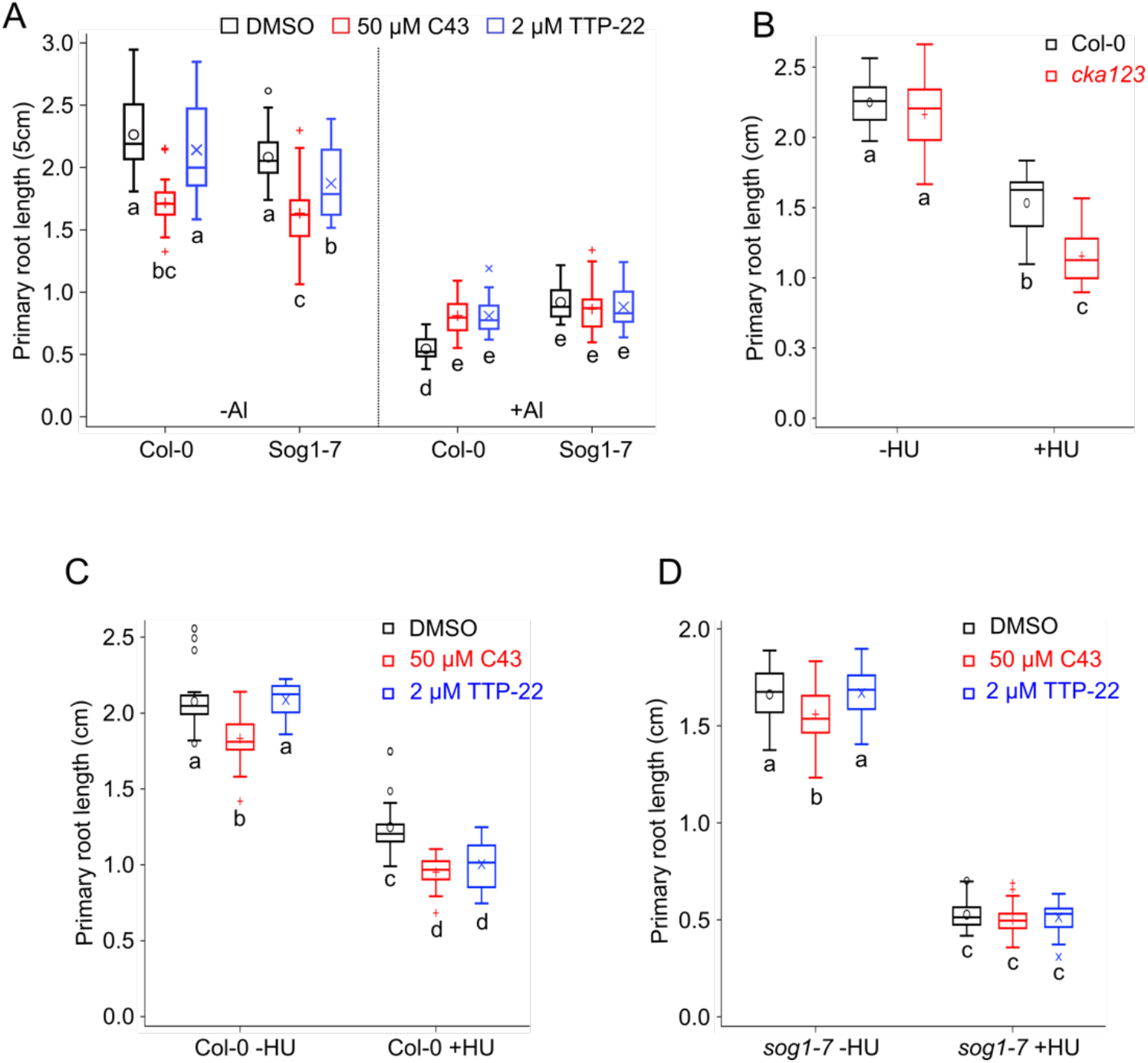
C43 and TTP-22 Operate Through SOG1. (A) Primary root length of 7-day-old wild-type (Col-0) and *sog1-7* plants grown in a soaked gel environment (pH 4.2) in the absence (-Al) or presence (+ Al) of 1.5 mM AlCl_3_, together with solvent control (DMSO), 50 μM C43 or 2 μM TTP-22. Data are represented by box and whisker plots. Significant differences (p < 0.01) as determined by mixed model analysis with Tukey’s post-hoc correction are indicated by letters below bars (n = 3, with > 8 plants in each repeat). (B) Primary root length of 7-day-old wild-type and *cka123* plants grown on 1/2 MS control medium (-HU) or supplemented with 1 mM HU (+HU). Data are represented by box and whisker plots. Significant differences (p < 0.01) as determined by mixed model analysis with Tukey’s post-hoc correction are indicated by letters below bars (n = 3, with > 8 plants in each repeat). (C) Primary root length of 7-day-old wild-type plants grown on 1/2 MS control medium (-HU) or supplemented with 1 mM HU (+HU), together with solvent control (DMSO), 50 μM C43 or 2 μM TTP-22. Data are represented by box and whisker plots. Significant differences (p < 0.01) as determined by mixed model analysis with Tukey’s post-hoc correction are indicated by letters below bars (n = 3, with > 8 plants in each repeat). (D) Primary root length of 7-day-old *sog1-7* plants grown on 1/2 MS control medium (-HU) or supplemented with 1 mM HU (+HU), together with solvent control (DMSO), 50 μM C43 or 2 μM TTP-22. Data are represented by box and whisker plots. Significant differences (p < 0.01) as determined by mixed model analysis with Tukey’s post-hoc correction are indicated by letters below bars (n = 3, with > 8 plants in each repeat).

SOG1 can be phosphorylated by both ATM and ATR (Yoshiyama et al., 2017) (Sjogren et al., 2015). ATM phosphorylation of SOG1 can occur at five different S/TQ sites upon treatment with DSB-inducing compounds such as zeocin. This phosphorylation is important for its activity, as demonstrated by the inability of a phospho-mutant allele to complement the *sog1* mutant. In addition to the ATM/ATR-dependent phosphorylation sites, 11 putative CK2 phosphorylation sites [T/SXXE/D, with S or T serving as the phospho-acceptor site (Meggio and Pinna, 2003)] are found within the full-length SOG1 protein. Among these, four sites are conserved among orthologous proteins (**Figure 6A**), stressing their potential importance for protein function. Of these, three cluster at the extreme C-terminus of SOG1, which is also the location of the ATM/ATR phosphorylation sites. To test putative phosphorylation of any of these sites, TAP-tagged SOG1 protein was extracted and purified from Arabidopsis cell cultures and subjected to phosphorylation analysis. Three phospho-peptides were detected (**Figure 6B** and **Supplemental Figure 4**). Among these, one corresponds to a predicted conserved CK2 consensus phosphorylation site (T423). To study the impact of phosphorylation on SOG1 activity, threonine at position 423 was changed to alanine (T423A). Both the mutant and wildtype *SOG1* allele were subsequently tested for their capability to complement the Al-resistant phenotype of *sog1-101* mutants (Takahashi et al., 2019). Two independent lines were selected for each, with equal transcription levels (**Supplemental Figure 5A**). Introducing the wild-type *SOG1* allele eliminated the Al resistance phenotype of the *sog1-101* mutant. In contrast, Al resistance was partially maintained for T423A lines, indicating that the mutation of T423 resulted in a partial loss of SOG1 activity (**Figures 7A,B and Supplemental Figure 5B**). Cotreatment of the T423A phospho-mutant plants with Al and C43 or TTP-22 did not result in a further increase in Al resistance compared to the treated wild type plants (both Col-0 and *sog1-101* mutants complemented with a wild-type *SOG1* allele) suggesting that under these experimental conditions T423 is the major CK2 phosphorylation site contribution to SOG1 activation (**Supplemental Figure 6**).

**Figure 6.**
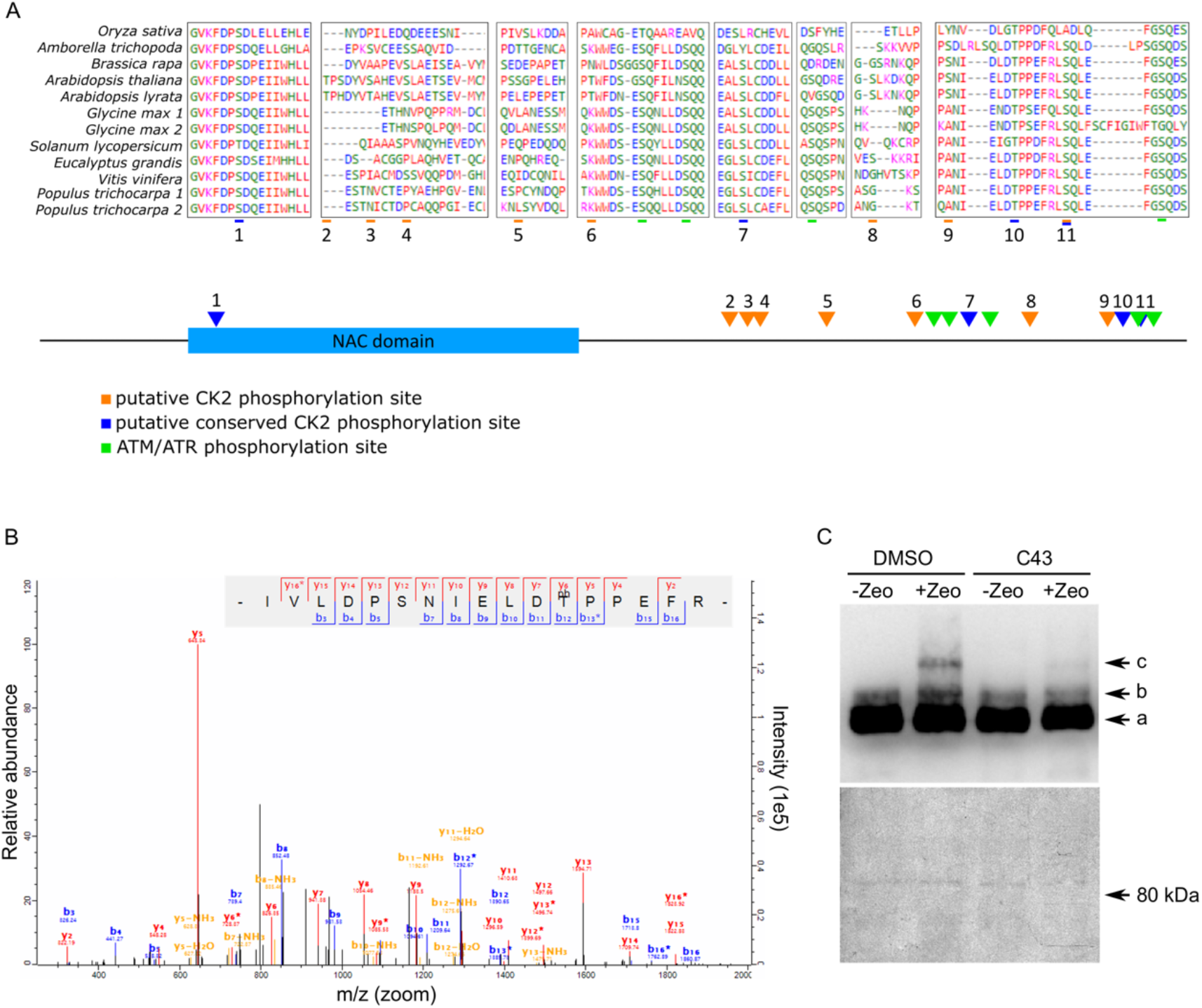
SOG1 Contains Several Evolutionary Conserved CK2 Phosphorylation Sites. (A) Putative CK2 phosphorylation sites were identified by searching for [ST]XXE occurrences in the *Arabidopsis* SOG1 protein. Protein sequences of SOG1 orthologs in different plant species were aligned using Clustal Omega and evolutionary conservation of the CK2 sites was determined. Blue and orange arrows indicate evolutionary conserved and non-conserved putative CK2 phosphorylation motifs, respectively, while green arrows indicate ATM/ATR phosphorylation motifs. (B) MS-MS fragment spectrum for the SOG1 412-428 tryptic fragment containing the T423 phosphorylation (phT). (C) Immunoblotting of total protein with anti-Myc antibody (upper panel). Total protein was extracted from 5-day-old SOG1:SOG1-Myc seedlings grown on MS medium with solvent control (DMSO) or 50 μM C43. Plants were transferred to liquid medium with (+) or without (-) 1 mM zeocin and total protein was extracted an hour later. The protein extracts were separated in an SDS–PAGE gel containing Phos-tag. Coomassie blue staining is displayed below. Non-phosphorylated, phosphorylated and hyperphosphorylated SOG1-Myc (bands a, b and c, respectively) are indicated by arrowheads.

The wild-type and mutant alleles were subsequently tested for their ability to induce cell death in vascular cells following treatment with the radiomimetic drug bleomycin. As previously reported (Furukawa et al., 2010; Johnson et al., 2018), *sog1* mutants failed to induce the cell death program upon induction of DSBs (**Supplemental Figure 5C**). This phenotype is rescued by reintroducing a wild-type *SOG1* allele, but was strongly attenuated in plants expressing the phospho-mutant allele (**Supplemental Figure 5C**), suggesting that SOG1 needs to be phosphorylated at T423 to be active under conditions that trigger DSBs. Strikingly, both phospho-mutant *SOG1* alleles rescued the HU hypersensitivity of the *sog1-101* allele to a level equal to that of the wild-type allele (**Figure 7C and Supplemental Figure 5D**), suggesting that under replication stress circumstances SOG1 activation occurs independently from T423 phosphorylation.

**Figure 7.**
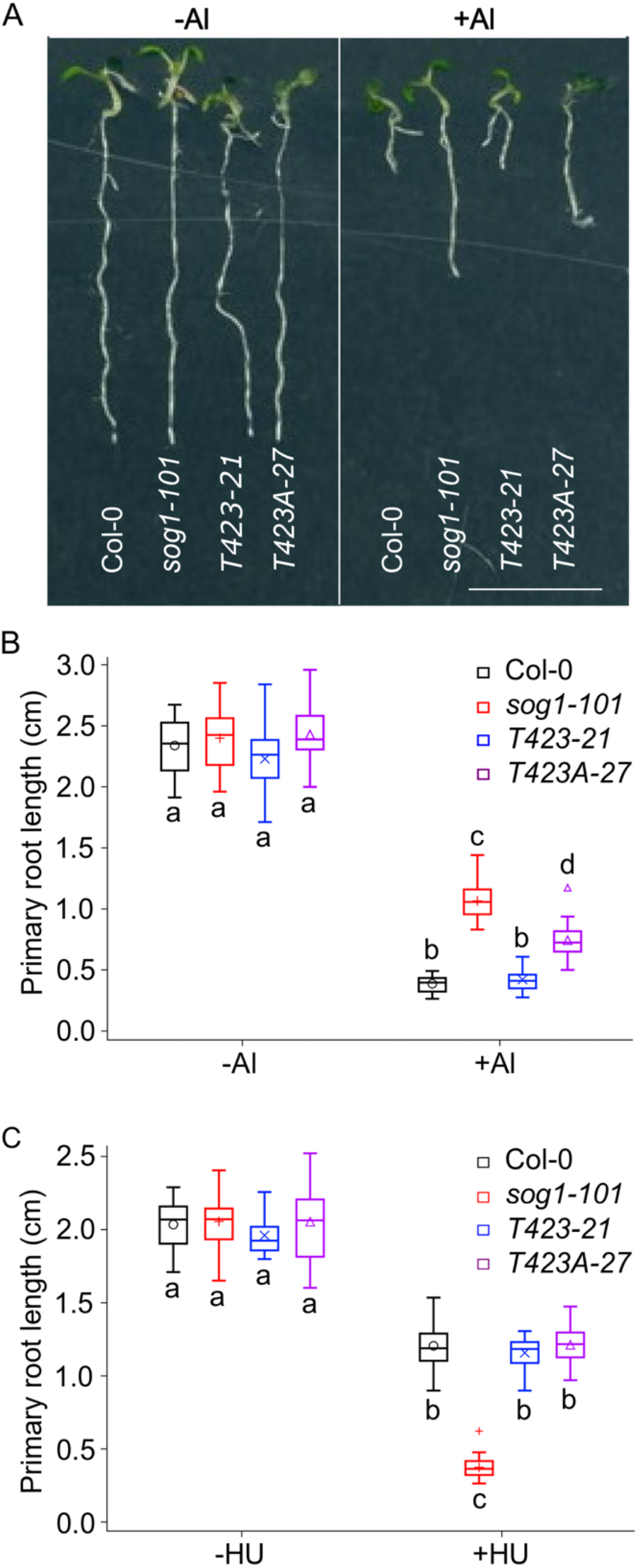
T423 Phosphorylation Is Required for SOG1 Activity Under Al Toxicity Conditions. (A) Wild-type (Col-0), *sog1-101* and *sog1-101* lines complemented with a wild-type (T423-21) or phospho-mutant (T423A-27) *SOG1* allele grown for 7 days in a soaked gel environment (pH 4.2) in the absence (-Al) or presence of 1.5 mM AlCl_3_. (+ Al). Plants were transferred to an agar plate to take pictures. Bar = 1 cm. (B) Quantification of data shown in (A). Data are represented by box and whisker plots. Significant differences (p < 0.01) as determined by mixed model analysis with Tukey’s post-hoc correction are indicated by letters (n = 3, with > 8 plants in each repeat). (C) Primary root length of wild-type, *sog1-101*, T423-21 and T423A-27 plants grown for 7 days on 1/2 MS control medium (-HU) or in the presence of 1 mM HU (+HU). Data are represented by box and whisker plots. Significant differences (p < 0.01) as determined by mixed model analysis with Tukey’s post-hoc correction are indicated by letters (n = 3, with > 8 plants in each repeat).

The phospho-mutant data suggested that SOG1 activity might be positively controlled through CK2 phosphorylation. To test this hypothesis, we studied the effects of CK2 inhibition through C43 or TTP-22 application on the SOG1-dependent transcriptional activation of DDR genes upon Al treatment. As reported by Sjogren et al. (2015), Al toxicity results in the transcriptional activation of both DNA repair (such as *RAD17* and *XRI1*) and cell cycle genes (such as *CYCB1;1*). This induction was partially suppressed through pretreatment of plants with either C43 or TTP-22 (**Supplemental Figure 7**). Additional evidence for CK2 operating upstream of SOG1 can be found in the observation that 21 out of 35 genes being induced in a CK2-dependent manner following γ-ray radiation (Moreno-Romero et al., 2012) represent SOG1 target genes (**Supplemental Figure 8**).

Following the induction of DSBs by the radiomimetic drug zeocin, SOG1 is targeted for phosphorylation by ATM (Yoshiyama et al., 2013). Therefore, it was tested whether CK2-dependent phosphorylation of SOG1 is required for its phosphorylation by ATM. As reported previously, treatment of seedlings for 24 h with zeocin resulted in the appearance of a slowly migrating band, previously demonstrated to correspond to the ATM-phosphorylated SOG1 isoform (Yoshiyama et al., 2013). Differently, when seedlings were pretreated with C43, the intensity of this band was strongly reduced, indicating that CK2-dependent phosphorylation of SOG1 is required for its phosphorylation by ATM in response to DSBs (**Figure 6C**).

### CK2 and SOG1 Deficiency Rescue Growth under Pi-Limiting Conditions

Response pathways to Al toxicity and low Pi availability converge on factors of Al tolerance. Loss of *ALUMINUM SESNSITIVE 3* (*ALS3*) not only results in hypersensitivity towards Al, but also results in an increased root growth inhibition under P-limiting conditions, due to an increased sensitivity to ROS that results from a greater bioavailability of Fe^3+^ (Larsen et al., 2005; Belal et al., 2015; Dong et al., 2017; Wang et al., 2019). Because of the reported correlation between Al toxicity and sensitivity to low P levels, it was tested whether C43 could rescue Arabidopsis root growth under Pi-deficient conditions. Initially, the low phosphate hypersensitive *als3-1* mutant background was used. Under high P conditions, no differences were observed in root length and meristem organization between control-of drug-treated *als3-1* mutant plants (**Figures 8A and 8B**). When grown on low P medium the *als3-1* mutants displayed a clear growth reduction (**Figure 8A**), as previously reported (Dong et al., 2017; Wang et al., 2019), and the root meristem was clearly disorganized (**Figure 8B**), being a sign of terminal differentiation and meristem exhaustion. In the presence of C43 or TTP-22, the *als3-1* root growth inhibition was slightly but significantly decreased (**Figure 8A**), with a clear recovery of root meristem organization (**Figure 8B**). Consistent with the hypothesis that CK2 controls SOG1 activity, *als3-1 sog1-7* roots showed partial suppression of the *als3-1* root growth phenotype and resulted in a recovery of root meristem organization (**Figures 8C and 8D**). Administration of either C43 or TTP-22 also resulted in an increased root growth under P-limiting conditions of wild-type plants (**Figure 8E**). Similarly, loss of SOG1 deficiency resulted in partial root growth rescue that could not be enhanced through application of C43 or TTP-22 (**Figure 8E**), suggesting that the root growth rescue by the drugs operates through the inhibition of SOG1 activity.

**Figure 8.**
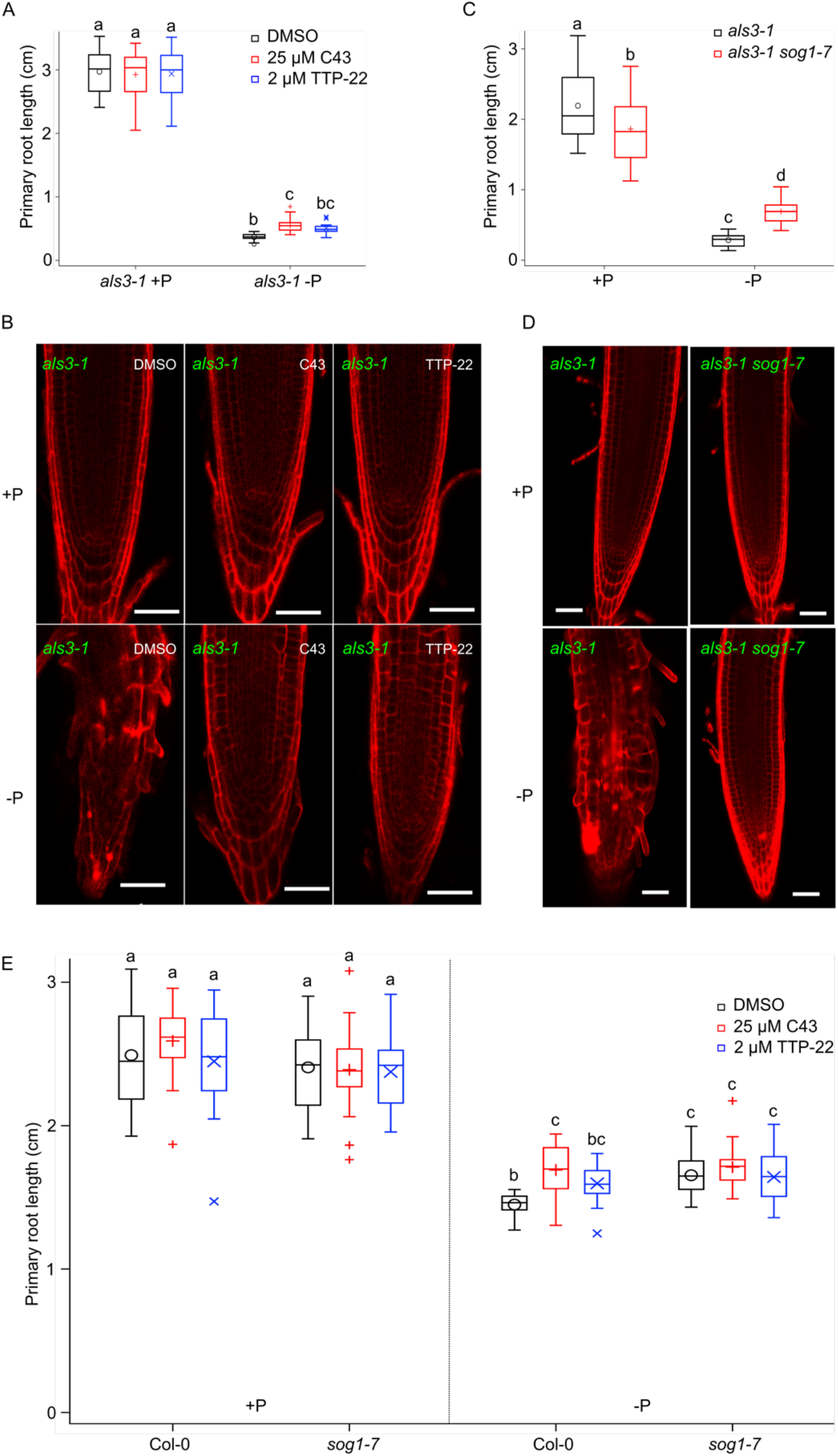
Root Growth and Meristem Organization under Pi-Limiting Growth Conditions Depend on CK2 and SOG1 Activity. (A) Primary root length of 7-day-old *als3-1* plants grown on P rich (1.25 mM, + P) or poor (10 μM, −P) medium together with solvent control (DMSO), 25 μM C43 or 2 μM TTP-22. Data are represented by box and whisker plots. Significant differences (p < 0.05) as determined by mixed model analysis with Tukey’s post-hoc correction are indicated by letters above bars (n = 3, with > 8 plants in each repeat). (B) Representative confocal microscopy images of plants shown in (A) stained with propidium iodide. Bar = 50 μm. (C) Primary root length of 7-day-old *als3-1* and *als3-1 sog1-7* plants grown on P rich (1.25 mM, + P) or poor (10 μM, −P) medium. Data are represented by box and whisker plots. Significant differences (p < 0.01) as determined by mixed model analysis with Tukey’s post-hoc correction are indicated by letters above bars (n = 3, with > 8 plants in each repeat). (D) Representative confocal microscopy images of plants shown in (C) stained with propidium iodide. Bar = 50 μm. (E) Primary root length of 7-day-old wild type (Col-0) and *sog1-7* plants grown on P rich (1.25 mM, + P) or poor (10 μM, −P) medium together with solvent control (DMSO), 25 μM C43 or 2 μM TTP-22. Data are represented by box and whisker plots. Significant differences (p < 0.05) as determined by mixed model analysis with Tukey’s post-hoc correction are indicated by letters above bars (n = 3, with > 8 plants in each repeat).

### C43 Application Results in Increased Pi Levels

As the knockout of rice CK2 activity been demonstrated to result in an intracellular increase of Pi (Chen et al., 2015) we similarly tested whether administration of C43 resulted in an increase of Pi uptake. In agreement with a CK2-inhibiting role of C43, drug application promoted a significantly increased Pi content (**Figure 9A**). ICP analysis indicated that C43 treatment mainly affected Pi homeostasis, as other chemical elements were not affected (**Supplemental Figure 9**). Pi content modifications are therefore not triggered by a major general effect of C43 on plant metabolism. In agreement with C43 being a CK2 inhibitor, the *cka123* mutant line displayed an increase in Pi accumulation compared to wild-type plants treated with C43 (**Figure 9A**). A minor increase in Pi content could be observed following treatment of *cka123* line with C43, suggesting that this *CK2* mutant still maintains some partial CK2 activity.

**Figure 9.**
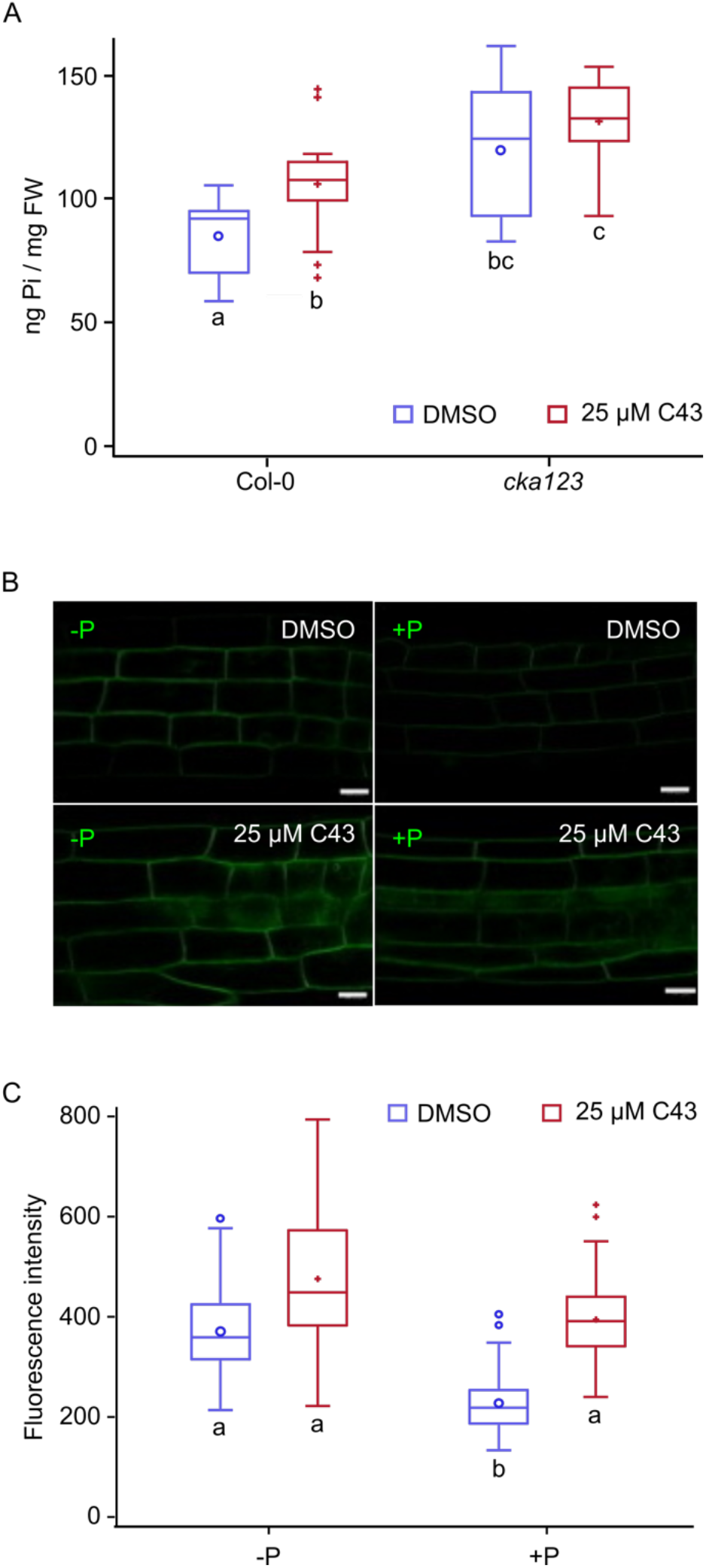
CK2 Controls Pi Homeostasis. (A) Soluble Pi level in roots of wild-type (Col-0) and *cka123* plants grown for two weeks on 1/10 MS medium supplemented with 100 μM P together with solvent control (DMSO) or 25 μM C43. Data are represented by box and whisker plots. Significant differences (p < 0.05) as determined by mixed model analysis with Tukey’s post-hoc correction are indicated by letters below bars (n = 5, with each sample containing 6 roots). (B) Subcellular localization of PHT1;1-GFP in root epidermal cells of 5-d-old seedlings grown in the absence (-P) or presence (+ P, 100 μM) of P and treated for 14 h with solvent control (DMSO) or 25 μM C43. (C) Quantification of GFP fluorescence for pictures shown in (B). Significant differences (p < 0.05) as determined by mixed model analysis with Tukey’s post-hoc correction are indicated by letters below bars (n = 5-8 with 20 measurements for each plant).

Previously, CK2 has been found to control Pi uptake through the control of the high-affinity Pi transporters PHT1 at the plasma membrane (Bayle et al., 2011; Chen et al., 2015). Under non-limiting Pi conditions, CK2 phosphorylates PHT1, by which it is triggered for destruction. Contrastingly, when Pi is scarce, PHT1 is not phosphorylated and can be seen to accumulate at the plasma membrane. To investigate the impact of C43 on PHT1 localization, we used seedlings containing the *p35S:PHT1;1-GFP* reporter construct (Bayle et al., 2011). The constitutive *Cauliflower Mosaic Virus 35S* promoter was used here to abolish strong transcriptional regulation affecting PHT1 accumulation in response to P treatment. Plants grown under low P conditions displayed a strong PHT1-GFP signal at the plasma membrane, irrespectively of whether they were control-treated or treated with 25 μM C43 (**Figure 9B**). As expected, under high P conditions, a decrease of the PHT1;1-GFP signal could be observed, matching previous data obtained for Arabidopsis and rice (Bayle et al., 2011; Chen et al., 2015). Contrastingly, a 14-h treatment with C43 resulted in PHT1;1-GFP accumulation to a level being equal to that observed under low P conditions (**Figure 9B**). Quantification of the GFP signals demonstrated that also under low P conditions, C43 administration resulted in a significant increase of the GFP signal (**Figure 9C**), again indicating that not all CK2 activity is inhibited, even at the low P concentration.

### Limiting Pi Levels Trigger an ATM-Dependent DDR Response

As loss of CK2 activity results in both an increase in Pi uptake and failure to activate SOG1, it was tested which of these two effects predominantly contributes to the rescue of the Pi limitation-induced growth phenotypes. First, it was tested whether the rescue of root growth of the *als3-1* mutant by C43 or TTP-22 was SOG1 dependent. Contrary to observed for the *als3-1* single mutant (**Figure 8A**), no root recovery could be detected in the *als3-1 sog1* double mutant upon addition of C43 or TTP-22 when growth in low P medium (**Figure 10A**), implying that the rescue of the Pi-limiting phenotype does operate via a pathway involving SOG1, and thus indicating that the C43/TTP-22 rescue phenotype depends on an attenuated DDR response. As additional evidence for this hypothesis, the involvement of ATR and ATM was analyzed by testing the response of *atm* and *atr* mutants under low P conditions. Under high P conditions, no significant difference in root growth could be observed for the both mutants (**Figures 10B and 10C**). Contrary, the *atm* mutant showed a minor but statistically significant better root growth under low P conditions (**Figure 10B**), whereas *atr* mutants displayed a response similar to that of wild-type plants (**Figure 10C**). These data demonstrate that ATM contributes to the growth arrest induced by Pi limitation.

**Figure 10.**
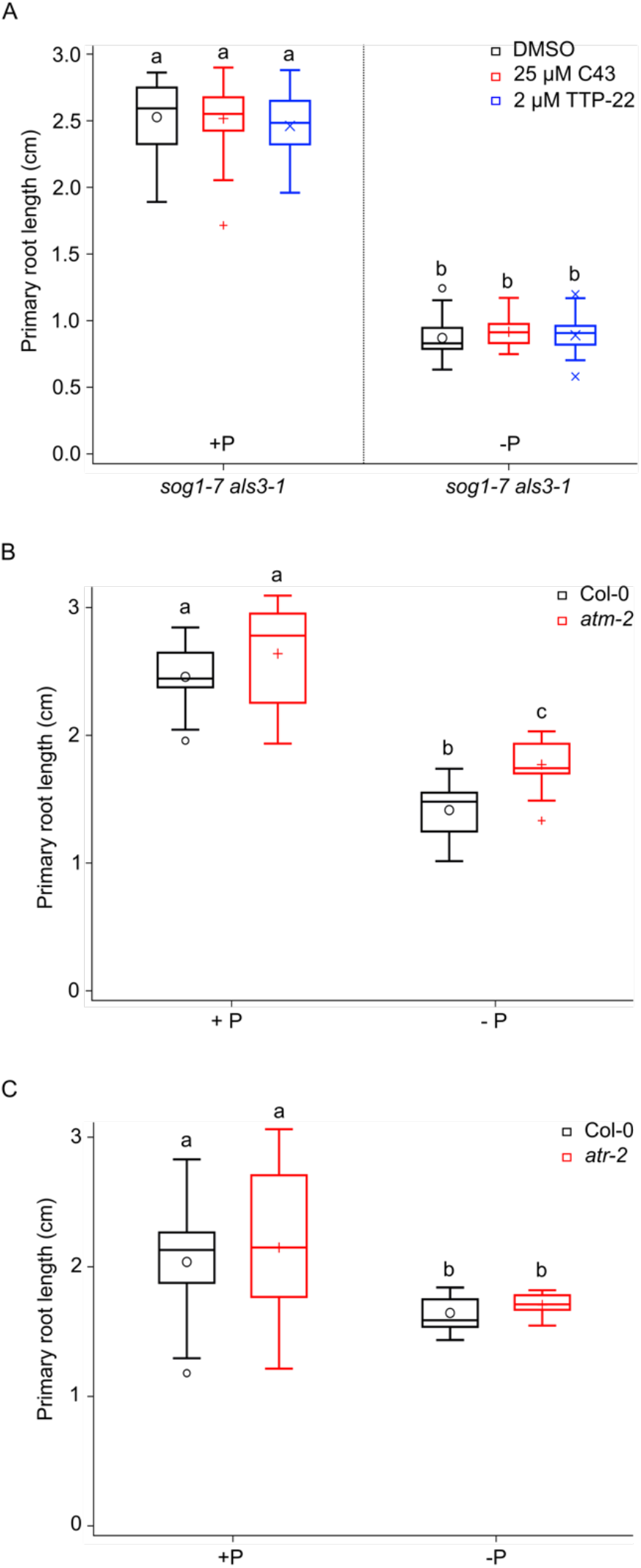
The Pi Limitation-Induced Growth Inhibition is ATM Dependent. (A) Primary root length of 7-day-old *als3-1 sog1-7* plants grown on P rich (1.25 mM, + P) or poor (10 μM, −P) medium together with solvent control (DMSO), 25 μM C43 or 2 μM TTP-22. Data are represented by box and whisker plots. Significant differences (p < 0.01) as determined by mixed model analysis with Tukey’s post-hoc correction are indicated by letters above bars (n = 3, with > 8 plants in each repeat). (B) Primary root length of 7-day-old wild-type (Col-0) and *atm-2* plants grown on P rich (1.25 mM, + P) or poor (10 μM, −P). Data are represented by box and whisker plots. Significant differences (p < 0.01) as determined by mixed model analysis with Tukey’s post-hoc correction are indicated by letters above bars (n = 3, with > 8 plants in each repeat). (C) Primary root length of 7-day-old wild-type and *atr-2* plants grown on P rich (1.25 mM, + P) or poor (10 μM, −P). Data are represented by box and whisker plots. Significant differences (p < 0.01) as determined by mixed model analysis with Tukey’s post-hoc correction are indicated by letters above bars (n = 3, with > 8 plants in each repeat).

## DISCUSSION

### CK2 Represents a Novel DDR Regulator in the Al Toxicity and Phosphate Limitation Response

CK2 represents a pleiotropic protein kinase with hundreds of potential substrates that are predominantly linked to signal transduction and gene expression (Pinna, 2002; Salinas et al., 2006). Here, we identified the plant CK2 as an important signaling component controlling root growth in response to Al toxicity and under conditions of Pi deficiency. Plant CK2 activity can be targeted through the chemical compounds TTP-22 and C43, the latter identified here through a chemical compound screen aimed at identifying chemicals that rescue plants from meristem exhaustion under otherwise toxic Al conditions. Although we could not demonstrate a biochemical activity for C43, probably due to its lower solubility and activity compared to TTP-22, genetic data strongly support an CK2 inhibitory role for both TTP-22 and C43. Not only does C43 share structural similarity to TTP-22, C43 mimics all TTP-22 induced phenotypes tested, including Al toxicity resistance, hypersensitivity to the replication inhibitor HU, and rescue from meristem exhaustion under P-limiting conditions. Additionally, the phenotypes of C43-treated plants are largely identical to those of untreated *CK2* knockdown plants, suggesting that CK2 is the only or main target of C43.

Our data indicate that CK2 contributes to the DDR response through phosphorylation of SOG1, fitting with reported hypersensitivity of a dominant-negative *CKA3* allele to γ-rays. UV-C and the DNA-alkylating agent methyl methanesulfonate (Moreno-Romero et al., 2012). A role for CK2 in cell cycle checkpoint control is supported by the effects of its inhibition in synchronized tobacco BY-2 cells, resulting in a premature condensation of DNA, implying lack of the G2/M checkpoint (Espunya et al., 1999). Additional proof for CK2 operating upstream of SOG1 can be found in the observation that 21 out of 35 genes being induced in a CK2-dependent manner following γ-ray radiation (Moreno-Romero et al., 2012) appear to be SOG1 target genes. Correspondingly, *CK2* knockdown mutant plants do not recover from exposure to 100 Gy and instead terminally differentiate, a phenotype similar to that of *sog1* mutant plants (Johnson et al., 2018), further supporting that CK2 and SOG1 operate in the same pathway.

We mapped the CK2 phosphorylation site within SOG1 to T423, being a highly conserved amino acid across SOG1 orthologous proteins, substantiating its importance. Mutation of T423 into a non-phosphorylated alanine residue only partly rescued the *sog1* mutant in terms of DNA damage-induced cell death and Al toxicity resistance, suggesting that CK2 phosphorylation is required to fully activate SOG1. At this moment, it is unknown how phosphorylation controls SOG1 activity. SOG1 is generally put forward as the plant’s ortholog of the mammalian p53 oncogene transcription factor (Yoshiyama, 2016). Strikingly, p53 is a well-characterized CK2 target, and interfering with its phosphorylation triggers susceptibility to UV-induced skin tumors (Bruins et al., 2004). CK2 phosphorylates p53 at the highly conserved S392 amino acid, promoting site-specific DNA-binding through stabilization of the p53 tetramer and tetramerization (Hupp et al., 1992; Sakaguchi et al., 1997; Keller and Lu, 2002), In contrast to the p53 protein that has been documented to be controlled at numerous levels (Dai and Gu, 2010), little is known about the posttranscriptional control of SOG1, apart from its activation through ATM/ATR-dependent phosphorylation. As SOG1 belongs to the family of NAC transcription factors, of which the members are known to operate as homo- or heterodimers, CK2 might control the dimerization status of SOG1 in a manner being similar to p53. When treated with either C43 or TTP-22, reduced SOG1 target gene activation was observed following Al treatment, suggesting possible reduced promoter binding affinity of the unphosphorylated SOG1 isoform to target genes, although currently such biochemical evidence is lacking. Strikingly, inhibition of CK2 activity interfered with ATM/ATR-dependent phosphorylation of SOG1 following treatment with zeocin. These data suggest a putative priming mechanism, where targets of ATM/ATR are only recognized after phosphorylation by CK2. As CK2 activity has been linked with many biological processes, such priming mechanism would allow to integrate the physiological condition of plants with DDR activation.

Strikingly, whereas the T423 residue is required for SOG1 activation under Al toxicity conditions, *sog1-8* mutant lines complemented with the phospho-mutant *SOG1* allele were as sensitive to HU as lines complemented with a wild type *SOG1* allele, suggesting that the T423 amino acid is not of importance for SOG1 activity under replication stress conditions. On the other hand, both wild type plants treated with C43 or TTP-22 as *cka123* mutants displayed a HU hypersensitivity that resembles that of *SOG1* mutant plants (Hu et al., 2015). It suggests that depending on the type of DNA damage different SOG1 residues might be targeted by CK2 phosphorylation. Alternatively, CK2 might control replication stress sensitivity through another checkpoint regulator.

### SOG1 and ATM Are Part of a Low Phosphate Level-Triggered Cell Cycle Checkpoint

Next to conferring Al toxicity resistance, both CK2 inhibition and SOG1 inactivation rescued the meristem exhaustion phenotype of *als3-1* mutant plants grow under Pi-limiting conditions. Likewise, the root growth inhibition phenotype was slightly but significantly rescued in a wildtype background by CK2 and SOG1 deficiency. P-limiting conditions affects root growth through a rapid inhibition of the elongation of cells within the root elongation zone and a sequential loss of cell proliferative activity of the stem cells. Both phenotypes depend on the accumulation of Fe^3+^. Whereas the phenotypes within the elongation zone have been attributed to malate-Fe^3+^-dependent accumulation of ROS and peroxidase activity that eventually affects cell wall stiffness (Balzergue et al., 2017; Wang et al., 2019), the meristem phenotypes have been credited to Fe^3+^-dependent accumulation of callose in the apoplast of the stem cell niche, interfering with the movement of non-cell autonomous stem cell regulators (Müller et al., 2015). The involvement of CK2 and SOG1 in the establishment of the P starvation phenotypes suggest that additionally to blocking intracellular transport, activation of a DDR-induced cell cycle arrest contributes to the exhaustion of the meristem. This is in particular clear in the *als3-1* mutant background. Although the exact substrate of the ALS3 transporter is yet to be determined, its structural similarity with the bacterial Fe exporter (Müller et al., 2015) and localization at the tonoplast (Dong et al., 2017) suggest a role for ALS3 in excess scavenging of reactive iron to the vacuole. We therefore speculate that in the *als3* mutants nuclear accumulation of Fe^3+^ might affect the DNA in a manner being similar to that of Al^3+^, resulting in DDR activation.

Next to the control of SOG1 activity, *Arabidopsis* CK2 was found to control Pi uptake, similarly as described for rice (Chen et al., 2015). Although rice CK2 activity has been reported to be repressed under P-limiting conditions (Chen et al., 2015), the observed rescue of both wild-type and *als3-1* mutant plants by CK2 inhibition under P-limiting conditions suggests that, at least in Arabidopsis, CK2 activity is not fully repressed under such conditions, allowing maintenance of SOG1 priming. Moreover, the observed rescue of root growth (for both wildtype and *als3-1* genotypes) and meristem organizing (for *als3-1)* by SOG1 deficiency in the absence of TTP-22 or C43 suggests that CK2’s role in the DDR damage response prevails in the induction of the root meristem Pi deficiency phenotypes. Possibly, CK2 operates in a tissuespecific manner. PHT1 activity has been predominantly attributed to non-dividing cell types including the root cap, root hairs and mature cortex cells (Mudge et al., 2002), whereas the DDR is of main importance in dividing cells. The model of a Pi limitation-activated cell cycle checkpoint is supported by the rescue root meristem phenotypes by the *atm* mutant. The identification of SOG1 and ATM as being part of a low-level Pi cell cycle checkpoint mechanism may therefore not only highlight an important novel physiological function for the plant DDR pathway, it might also hint towards the development of novel approaches to cope with the agricultural problem of Pi limitation.

## METHODS

### Plant Materials and Growth Conditions

Unless stated otherwise, Arabidopsis plants were grown under long-day conditions (16 h of light/8 h of darkness, Lumilux Cool White lm, 50 to 70 μmol m^−2^ s^−1^) at 22°C on half-strength Murashige and Skoog (MS) germination medium (Murashige and Skoog, 1962), 10 g/L sucrose, 0.5 g/L MES, pH 5.7 and 10 g/L plant tissue culture agar (Neogen). The *cka123* mutant (N67786) was acquired from the Nottingham Arabidopsis Stock Centre (NASC). The *sog1-7, sog1-101, als3-1, als3-1 sog1-7, atm-2* and *atr-2* have been previously described (Nezames et al., 2012; Sjogren et al., 2015).

### Chemical Screen

Col-0 plants that contain the *QC25:GUS* reporter were used for a compound screen. Arabidopsis seeds were surface-sterilized and incubated for 2 d at 4°C. Then seeds were sown into 96-well plates (Thermo Fisher Scientific, MA, USA) with 150 μl per well. The medium (pH 4.2) consisting of 1 mM KNO_3_, 0.2 mM KH_2_PO_4_, 2 mM MgSO_4_, 0.25 mM (NH_4_)_2_SO_4_, 1 mM Ca(NO_3_)_2_, 1 mM CaSO_4_, 1 mM K_2_SO_4_, 1 μM MnSO_4_, 5 μM H_3_BO_3_, 0.05 μM CuSO_4_, 0.2 μM ZnSO_4_, 0.02 μM NaMoO_4_, 0.1 μM CaCl_2_, 0.001 μM CoCl_2_ and 1% sucrose. Three days after germination, the control nutrient medium was replaced with 150 μl per well Al-containing medium (medium with additional 0.5 mM AlCl_3_, pH 4.2). Subsequently, compounds of the Pharmacological Diversity Set (Enamine) were added with a Te-MO robot (Tecan, Switzerland) at a final concentration of 50 μM. DMSO-treated samples without Al were used as positive controls and DMSO-treated samples with Al were used as negative controls. After 5 d of growth, overnight GUS staining was performed, after which the seedlings were washed 2 to 3 times for 10 min with phosphate buffer (pH 7.2). Samples were mounted in lactic acid and photographed with a stereomicroscope (Olympus BX51 microscope).

### Al Treatment

Arabidopsis Col-0 wild-type and mutant seeds were surface-sterilized and cold-stratified at 4°C for 2 d in the dark. Then seeds were sown on either soaked gel plates or hydroponic plates as previously described (Nezames et al., 2012). For soaked gel plates, 0.125% Gellan gum (Gell-8Gro; ICN Biomedicals, CA, USA) was added to the control nutrient medium. Then the solidified nutrient medium was soaked with 20 ml of nutrient medium ± 1.5 mM AlCl_3_ (pH 4.2) for 2 d, after which the soaking solution was removed, and seeds were plated and grown for 7 d.

### Phosphate Treatment

Seeds were surface-sterilized, cold-stratified at 4°C for 2 d in the dark and germinated on 0.8% (w/v) washed agar medium (pH 5.5) containing 5 mMKNO_3_, 2.5 mM KH_2_PO_4_, 2 mM MgSO_4_, 2 mM Ca(NO_3_)_2_, 50 μM Fe-EDTA. 70 μM H_3_BO_3_, 14 μM MnCl_2_, 0.5 μmCuSO_4_, 1 μM ZnSO_4_, 0.2 μM NaMoO_4_, 10 μM NaCl, and 0.01 μM CoCl_2_ and 0.5% (w/v) sucrose. For low or -Pi medium, 10 μM KH_2_PO_4_ was used. For agar washing, per 100 g plant tissue culture agar (Neogen) was washed with 5 L Milli-Q water for 4 times and then incubated with dialysis tubing (INTERCHIM, T4-25-30) filled with Dowex (Sigma-Aldrich, 14015-U) for 12 h. After that, the supernatant was discarded, and the agar was dried on the plastic tray at 60°C for at least 2 days.

### GUS Assays

GUS staining experiments were conducted as previously described (Beeckman and Engler, 1994). After overnight staining, samples were mounted in lactic acid and imaged with an Olympus BX51 stereomicroscope.

### Generation of *SOG1* Complementation Lines

To generate the T423 transgene lines, the full *SOG1* genomic ORFs without stop codon were cloned into the pDONR221 (Invitrogen, CA, USA) vector and the 1,990-bp promoter region into the pDONR-P4P1 vector using Invitrogen BP-clonase according to manufacturer’s instructions. Endogenous promoter-driven translational fusions were created with the MultiSite Gateway technology that combined the endogenous promoter and 3xMyc fragments downstream of open reading frames in the pH7m34GW destination vector (Karimi et al., 2007). To make the T423A mutant lines, the threonine at position 423 was changed to alanine by site-directed mutagenesis by PCR (Zheng et al., 2004). The destination constructs were transformed into *sog1-101* plants by floral dipping (Clough and Bent, 1998).

### Morin Staining

Arabidopsis seedlings were grown in control nutrient medium for 5 d. Then plants were exposed to AlCl_3_ for 1 h. Morin staining was conducted as previously described (Larsen et al., 1996). Briefly, seedlings were washed in Mes buffer (pH 5.5) for 10 min and stained with 100 μM Morin (Sigma-Aldrich, MO, USA) in the same buffer for 1 h. Morin fluorescence was visualized using a Zeiss LSM 710 microscope.

### Callose Staining

Five-d-old seedlings were exposed to 50 μM AlCl_3_ for 24 h, after which plants were fixed with 10% formaldehyde, 5% glacial acetic acid, and 45% ethanol and vacuum-infiltrated for 4 h. Then seedlings were incubated in 0.1% aniline blue (pH 9.0, 0.1 M K_3_PO_4_). Callose production was visualized using a Zeiss LSM 710 microscope.

### Real-Time PCR Analysis

RNA was extracted from Arabidopsis tissues with the RNeasy mini kit (Qiagen, Germany). After DNase treatment with the RQ1 RNase-Free DNase (Promega, WI, USA), cDNA was synthesized with the iScript cDNA synthesis kit (Bio-Rad, CA, USA). Quantitative RT-PCR was performed using the SYBR Green kit (Roche, Belgium) with 100 nM primers and 0.125 μl of RT reaction product in a total volume of 5 μl per reaction. Reactions were run and analyzed on the LightCycler 480 (Roche, Belgium) according to the manufacturer’s instructions. Each reaction was done in three technical and three biological repeats. Expression levels were normalized to expression of the *ACTIN2* and *EF-1a* reference genes. Fold changes were calculated using the 2^-ΔΔCt^ method. Primers used are given in Supplemental Table 1.

### Kinase assays

Kinase assays were carried out as described in (Dennis et al., 2009). Briefly, 1 pmol of CK2 kinase was incubated with 30 pmol HD2B (Substrates kindly provided by Prof. Karen Browning) with 200 cpm/pmol γ-32P ATP in 50 mM Tris-HCl, pH 7.6, 5 mM MgCl2 2.4 mM DTT in presence or absence of compound dissolved in DMSO. Reaction mixtures were incubated for 30 min at 37°C. The reaction was stopped by adding SDS-PAGE loading dye and boiling the samples for 5 min at 95°C. Samples were separated by SDS-PAGE analysis prior to autoradiography.

### Thermal Shift Assays

The compounds (TTP-22 and C43) were dissolved in DMSO to a stock concentration of 50 mM. Reaction mixtures of 25 μl total volume consisting of 5 μl CK2 (4 μM), 1x Sypro Orange and either DMSO, TTP-22 or C43 and 25 mM Tris pH 7.5 150 mM NaCl were loaded in Hardshell 96-well PCR plates (BioRad, Hercules, USA). Triplicates of every condition were prepared. Plates were sealed with Optical-Quality Sealing Tape (BioRad, Hercules, USA) and thermal denaturation was performed in a CKF Connect Real-Time System. The samples were subject to a gradient slope from 10°C to 95°C with a ramp rate of 0.025 °C/min.

### SOG1 Phosphorylation Assay

Five-day-old *ProSOG1:SOG1-Myc* (Yoshiyama et al., 2013) seedlings were transferred to MS liquid medium containing 0 or 1 mM zeocin. After a 1-h incubation, a pool of root tips (from ~100 seedlings) was excised and ground in the following buffer: 10 mM Tris (pH 7.6), 150 mM NaCl, 2 mM EDTA, 0.5% (v/v) Nonidet P-40 (Nacalai Tesque), 1 mM DTT, protease inhibitor cocktail (Sigma-Aldrich), and a phosphatase inhibitor cocktail (0.1 mM Na3VO4, 1 mM NaF, 60 mM b-glycerophosphatase, and 20 mM p-nitrophenylphosphate). The slurry was centrifuged (20,000g) twice to remove debris, and the supernatant was recovered and used for subsequent analysis. Proteins (1 μg) were separated on an 8% SDS-PAGE gel containing 40 μM MnCl_2_ and 20 μM Phos-tag (NARD Institute), which was used for identification of phosphorylated SOG1 protein. Phosphorylated SOG1 is visualized as bands that migrate more slowly than those of nonphosphorylated proteins. After electrophoresis, the proteins were electroblotted to a PVDF membrane (Merck Millipore) in the following buffer: 6.3 mM NaHCO3, 4.3 mM Na2CO3, pH 9.5, and 20% methanol. Because SOG1-Myc can be detected using an anti-Myc antibody, the membrane was incubated for 2 h at room temperature in the anti-Myc primary antibody A-14 (1:2000 dilution, lot no. F0810; Santa Cruz Biotechnology), rinsed three times with 1X TBST (0.01 M Tris-HCl, 0.15 M NaCl, and 0.05% Triton X-100), and incubated with an anti-rabbit immunoglobulin horseradish peroxidase-conjugated secondary antibody (1:4000 dilution, lot no. 187662; Promega) to detect SOG1-Myc. Next, the membrane was washed three times with 1X TBST and processed with a LAS-4000 luminescent image analyzer (Fujifilm) after incubation with the ECL Prime enhanced chemiluminescence kit (GE Healthcare).

### Determination of SOG1 phosphosites

Cloning of transgene encoding N-terminal GS^rhino^ tag SOG1 fusion under control of the constitutive *cauliflower tobacco mosaic virus 35S* promoter, transformation of Arabidopsis cell suspension cultures (PSB-D) with direct selection in liquid medium and TAP experiments were performed with 100 mg of total protein extract as input as described in (Van Leene et al., 2015). Bound proteins were digested on-bead after a final wash with 500 μL 50 mM NH_4_HCO_3_ (pH 8.0). Beads were incubated with 1 μg Trypsin/Lys-C in 50 μL 50 mM NH_4_OH and incubated at 37°C for 4 h in a thermomixer at 800 rpm. Next, the digest was separated from the beads, an extra 0.5 μg Trypsin/Lys-C was added and the digest was further incubated overnight at 37°C. Finally, the digest was centrifuged at 20800 rcf in an Eppendorf centrifuge for 5 min, the supernatant was transferred to a new 1.5 mL Eppendorf tube, and the peptides were dried in a Speedvac and stored at −20°C until MS analysis.

The obtained peptide mixtures were introduced into an LC-MS/MS system, the Ultimate 3000 RSLC nano (Dionex, Amsterdam, The Netherlands) in-line connected to an Orbitrap Elite™ Hybrid Ion Trap-Orbitrap Mass Spectrometer (Thermo Fisher Scientific, Bremen, Germany). The sample mixture was loaded on a trapping column (made in-house, 100 μm internal diameter (I.D.) x 20 mm (length), 5 μm C18 Reprosil-HD beads, Dr. Maisch GmbH, Ammerbuch-Entringen, Germany). After back-flushing from the trapping column, the sample was loaded on a reverse-phase column (made in-house, 75 μm I.D. x 150 mm, 5 μm C18 Reprosil-HD beads, Dr. Maisch). Peptides were loaded with solvent A (0.1% trifluoroacetic acid, 2% acetonitrile), and separated with a linear gradient from 98% solvent A’ (0.1% formic acid) to 50% solvent B’ (0.1% formic acid and 80% acetonitrile) at a flow rate of 300 nl/min, followed by a wash step reaching 100% solvent B’. The mass spectrometer was operated in data-dependent, positive ionization mode, automatically switching between MS and MS/MS acquisition for the 20 most abundant peaks in a given MS spectrum. In the Orbitrap Elite, full scan MS spectra were acquired in the Orbitrap at a target value of 3E6 with a resolution of 60,000. The 20 most intense ions were then isolated for fragmentation in the linear ion trap, with a dynamic exclusion of 20 seconds. Peptides were fragmented after filling the ion trap at a target value of 3E4 ion counts.

The Thermo raw files were processed using the MaxQuant software (version 1.6.4.0). Data were searched with the built-in Andromeda search engine against the TAIR10 database (TAIR10_pep_20101214.fasta). Variable modifications were set to oxidation of methionines, N-acetylation of protein N-termini and phosphorylation of serines, threonines, tyrosines. Mass tolerance on precursor (peptide) ions was set to 4.5 ppm and on fragment ions to 0.5 Da. Enzyme was set to Trypsin/P, allowing for 2 missed cleavages, and cleavage was allowed when arginine or lysine was followed by proline. PSM, protein and site identifications were filtered using a target-decoy approach at false discovery rate (FDR) of 1%.

The mass spectrometry proteomics data have been deposited to the ProteomeXchange Consortium (http://proteomecentral.proteomexchange.org) via the PRIDE partner repository (Perez-Riverol et al., 2019) with the dataset identifier PXD018727 (Reviewer account details: **Username:**; **Password:** ETGyIfhC).

### Inorganic Pi, Total P and Ion Content

Pi content was calculated as previously described (Misson et al., 2004) from rosettes and roots of in vitro-grown plantlets. Total phosphate and ions content was analyzed by ICP OES as previously described (Hirsch et al., 2006). Briefly plantlets were grown on medium holding 1/10th Murashige and Skoog germination medium (Murashige and Skoog, 1962), 0.5% sucrose, 2 μM FeCl_2_, 100 μM KH_2_PO_4_ and 8 g/l agar (Sigma). In order to avoid contamination by the medium, care was taken to collect only plants that did not enter agar and samples were rinsed with distilled water before analysis.

### Confocal Imaging and Quantification of GFP Fluorescence

The *p35S:PHT1;1-GFP* line was analyzed by confocal laser scanning microscopy using a Zeiss LSM780 inverted confocal microscope equipped with argon ion (488 nm) and He-Ne lasers. A C-Apo Corr FCS water immersion objective (numerical aperture = 1.2; ref. 4217679970711) was used. For quantification experiments, eight plantlets were observed per condition. Fluorescence from 20 regions of interest per plant was quantified using ImageJ software (http://rsb.info.nih.gov/ij/download.html). The quantification of GFP fluorescence was performed as previously described (Balzergue et al., 2017), with the following modifications: a 4 images Z-stack (separated by 2-μm distance) was used to study the membranes of the root. Images were acquired in 12 bits using Zen black software (Zen black 2012 SP2 Version 11.0), then converted to a maximal projection image. A 120 μm X 60 μm rectangle was defined at 200 μm from the root tip (roughly in the transition zone), in which a region of interest (6 μm X 6 μm) inside each nucleus was defined for the measurement of fluorescence.

### CKA1 Co-Crystallization Studies

Expression and purification of AtCKA1 was performed as previously described (Demulder et al., 2020). The protein was concentrated to 4 mg/ml in 25 mM Tris pH 7.5, 150 mM NaCl, 1 mM TCEP. TTP-22 or C43 were dissolved in DMSO at a concentration of 10 mM and added to the concentrated protein in a compound:protein ratio of 1:9. The mixture was incubated for half an hour at room temperature. Soluble protein was separated from aggregates by centrifugation for 5 min at 14,000 rpm in an Eppendorf centrifuge. Hanging drop crystallization screens were set up manually with drops consisting of 1 μl of protein solution and 1 μl of precipitant solution equilibrated against 300 μl of precipitant solution at 20°C. Plates were set up using the Hampton Research and Pact Premier screens. The crystals for the TTP-22 complex were obtained in 0.2 M sodium citrate tribasic dehydrate, 0.1 M Bis-Tris propane pH 8.5, 20% w/v PEG 3350. A crystal was cryo-protected by increasing the PEG 3350 concentration to 30% prior to flash-freezing in liquid nitrogen. X-ray data were collected at 100 K at beamlines Proxima-1 of the SOLEIL synchrotron source (Saint-Aubin, France). The data were indexed, scaled and merged with XDS (Kabsch, 2010) using the xdsme interface (https://github.com/legrandp/xdsme). The structure was determined by molecular replacement using the co-ordinates of AtCKA1 (PDB entry 6XX6) stripped of water molecules as search model using Phaser. This solution was further refined using Phenix. refine (Liebschner et al., 2019) and rebuilt using Coot (Emsley et al., 2010). A maximum likelihood target was used throughout. The final refinement cycles included TLS refinement (one TLS group per chain).

### Statistical Analysis

Statistical analyses were done as indicated in the figure legends, using the Mixed Model procedure in the SAS Studio software with Tukey-correction for multiple testing. Details for each experiment can be found in Supplemental Data Set 1.

## Supplemental Data

The following materials are available in the online version of this article.

**Supplemental Figure 1.** Dose-dependent Rescue of Primary Root Growth by C43 and TTP-22 Under Al Toxic Conditions.
**Supplemental Figure 2.** Callose and Al Accumulation in TTP-22-Treated Plants
**Supplemental Figure 3.** TTP-22 Binds and Inhibits CKA1
**Supplemental Figure 4.** SOG1 phosphorylation sites
**Supplemental Figure 5** Molecular and Phenotypic Characterization *sog1-101* Complementation Lines
**Supplemental Figure 6.** T423 represents a major CK2 phosphorylation site contributing to SOG1 activation.
**Supplemental Figure 7.** C43 and TTP-22 Repress Expression of SOG1 Target Genes
**Supplemental Figure 8**. CK2-Dependent Upregulation of Genes Shows Statistically Significant Enrichment for SOG1 targets
**Supplemental Figure 9.** C43 Does Not Affect Mineral Composition of Roots
**Supplemental Table 1.** List of primer sequences
**Supplemental Data Set 1.** Statistical analysis.

## ACKNOWLEDGMENTS

The authors thank Annick Bleys for help in preparing the manuscript, the VIB proteomics core for LC-MSMS analysis, Christian Godon for assistance to use ImageJ software to quantify GFP, and Hilde van Den Daele for making figures. This work was supported by grants of the Research Foundation Flanders (G011420N) and ERA-NET for Coordinating Plant Sciences (Al-UCIDATE). M.D. indebted to the Research Foundation Flanders for a predoctoral fellowship.

## AUTHOR CONTRIBUTIONS

P.W., M.D., P.D., T.E., K.O.Y, I.V., M.G., D.E. and T.D. conducted the experiments; G.D.J., P.L, D.A., L.N., T.D., R.L. and L.D.V. designed the experiments and P.W., M.D., R.L. and L.D.V. wrote the paper.

---

**Supplemental Figure 1.**
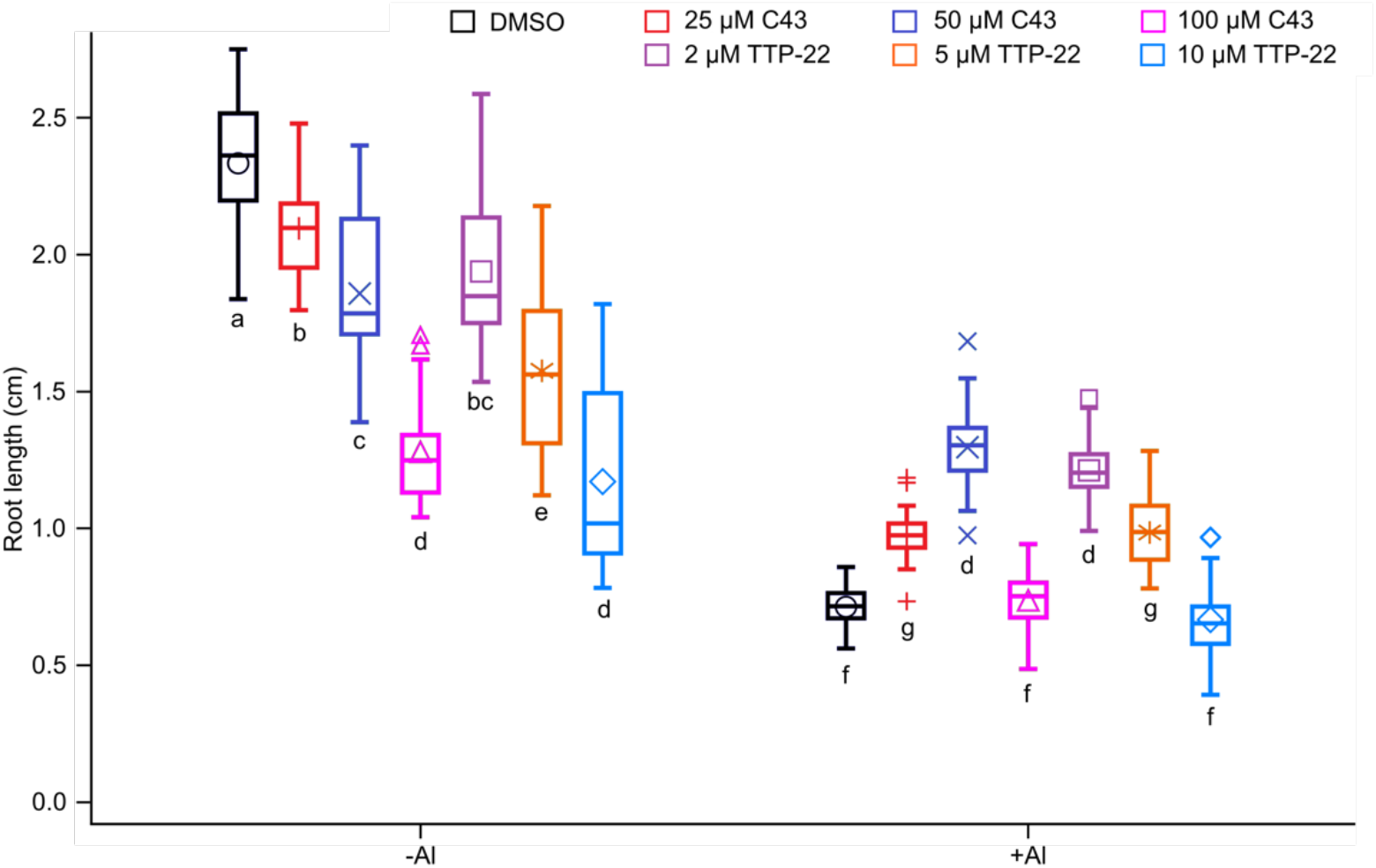
Dose-dependent Rescue of Primary Root Growth by C43 and TTP-22 Under Al Toxic Conditions (Supports Figures 2 and 3). Primary root length of 7-day-old wild-type (Col-0) seedlings grown in a soaked gel environment (pH 4.2) in the absence (-Al) or presence (+ Al) of 1.5 mM AlCl_3_, together with solvent control (DMSO) or indicated concentration of C43 or TTP-22. Data are represented by box and whisker plots. Significant differences (p < 0.05) as determined by mixed model analysis with Tukey’s post-hoc correction are indicated by letters above bars (n = 3, with > 8 plants in each repeat).

**Supplemental Figure 2.**
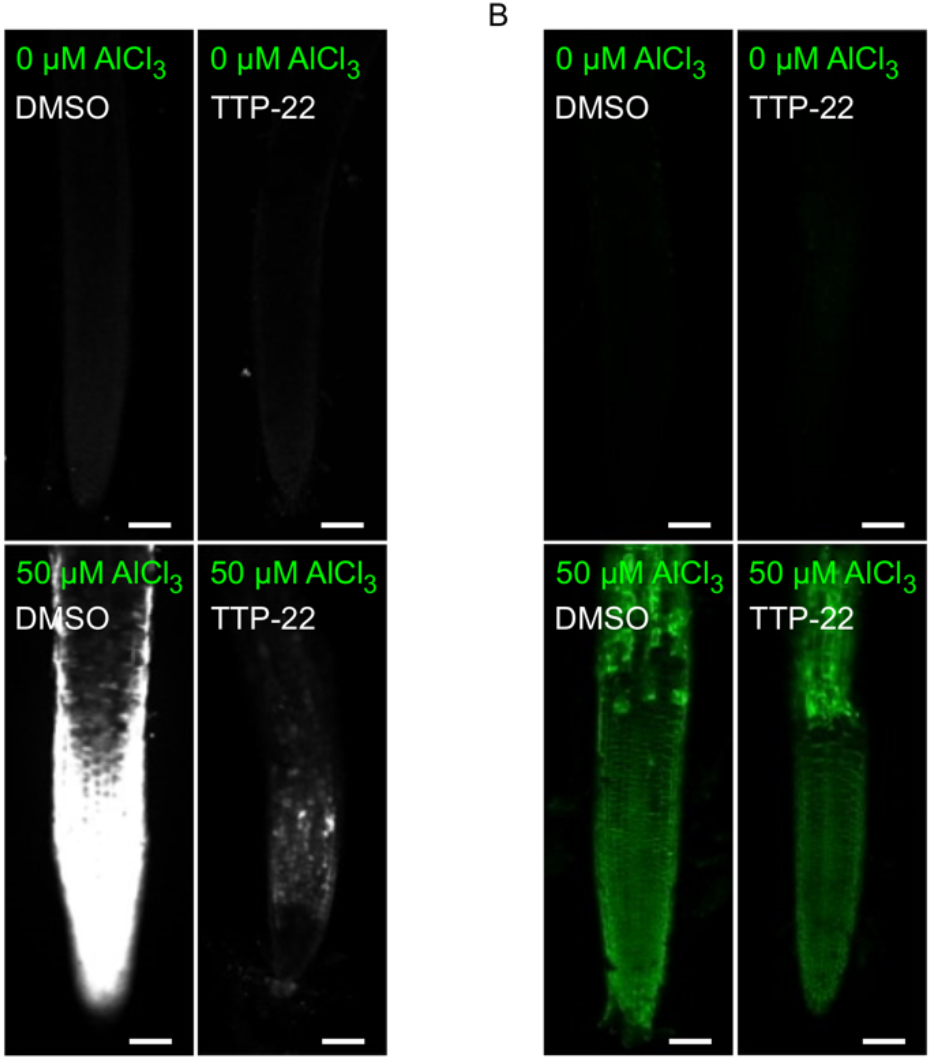
Callose and Al Accumulation in TTP-22-Treated Plants (Supports Figure 3) (A) Five-day-old wild-type (Col-0) seedlings hydroponically grown (at pH 4.2) in the presence of solvent control (DMSO) or 5 μM TTP-22 and exposed to either 0 (control) or 50 μM AlCl_3_ for 24 h. Seedling were stained with aniline blue and visualized using fluorescence microscopy for callose deposition. Bars = 50 μm. (B) Five-day-old wild-type seedlings hydroponically grown (at pH 4.2) in the presence of solvent control (DMSO) or 5 μM TTP-22 and exposed to either 0 (control) or 50 μM AlCl_3_ for 1 h. Seedlings were Morin stained and visualized using fluorescence microscopy for Al accumulation. Bars = 50 μm.

**Supplemental Figure 3.**
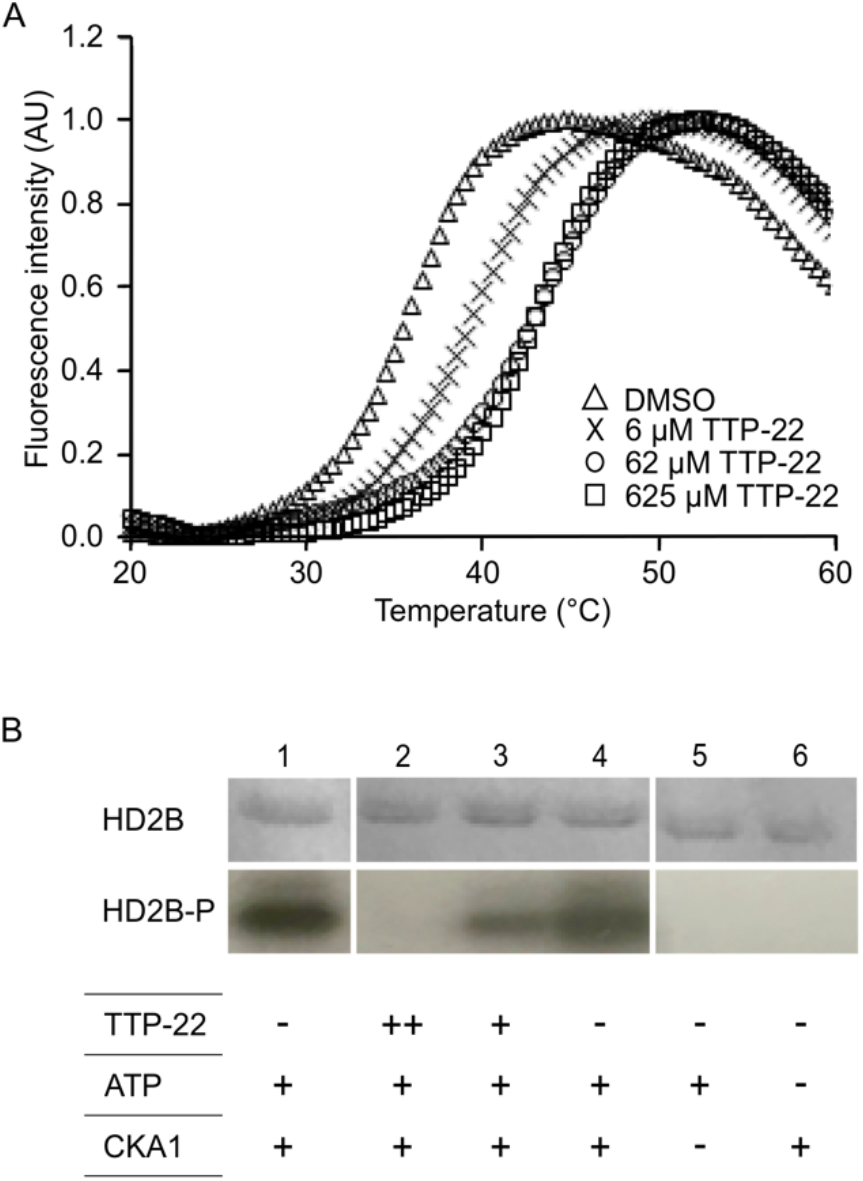
TTP-22 Binds and Inhibits CKA1 (Supports Figure 3 and 4) (A) CKA1 protein was subjected to a fluorescent thermal shift assay to monitor thermal stabilization in the presence of different concentrations of TTP-22 or incubated with DMSO. An upward shift of the melting temperature indicates that the compound binds and enhances the thermal stability of CK2A1. The average values of triplicates are displayed. Standard errors did no exceed 15% of the mean. (B) HD2B substrate (30 pmol) was incubated with CKA1 (1 pmol) in the presence of γ −^32^P ATP with or without de addition of TTP-22 (++ = 625 μM and + = 12 μM). Reaction mixtures were separated by SDS-PAGE and analyzed by autoradiography.

**Supplemental Figure 4.**
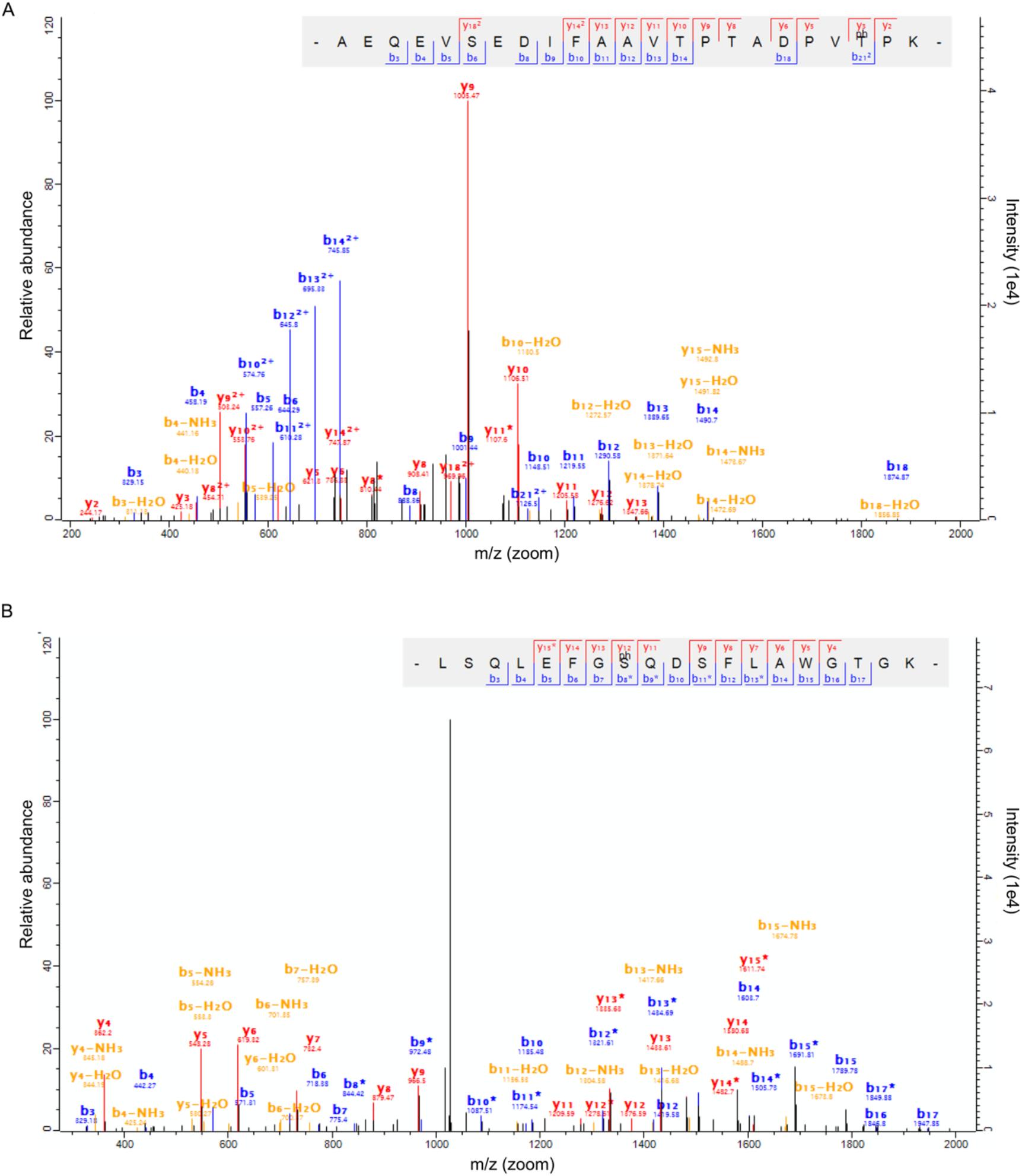
SOG1 phosphorylation sites (Supports Figure 6) (A) MS-MS fragment spectrum for the SOG1 224-246 tryptic fragment containing the T244 phosphorylation. (B) MS-MS fragment spectrum for the SOG1 429-447 tryptic fragment containing the S436 phosphorylation.

**Supplemental Figure 5.**
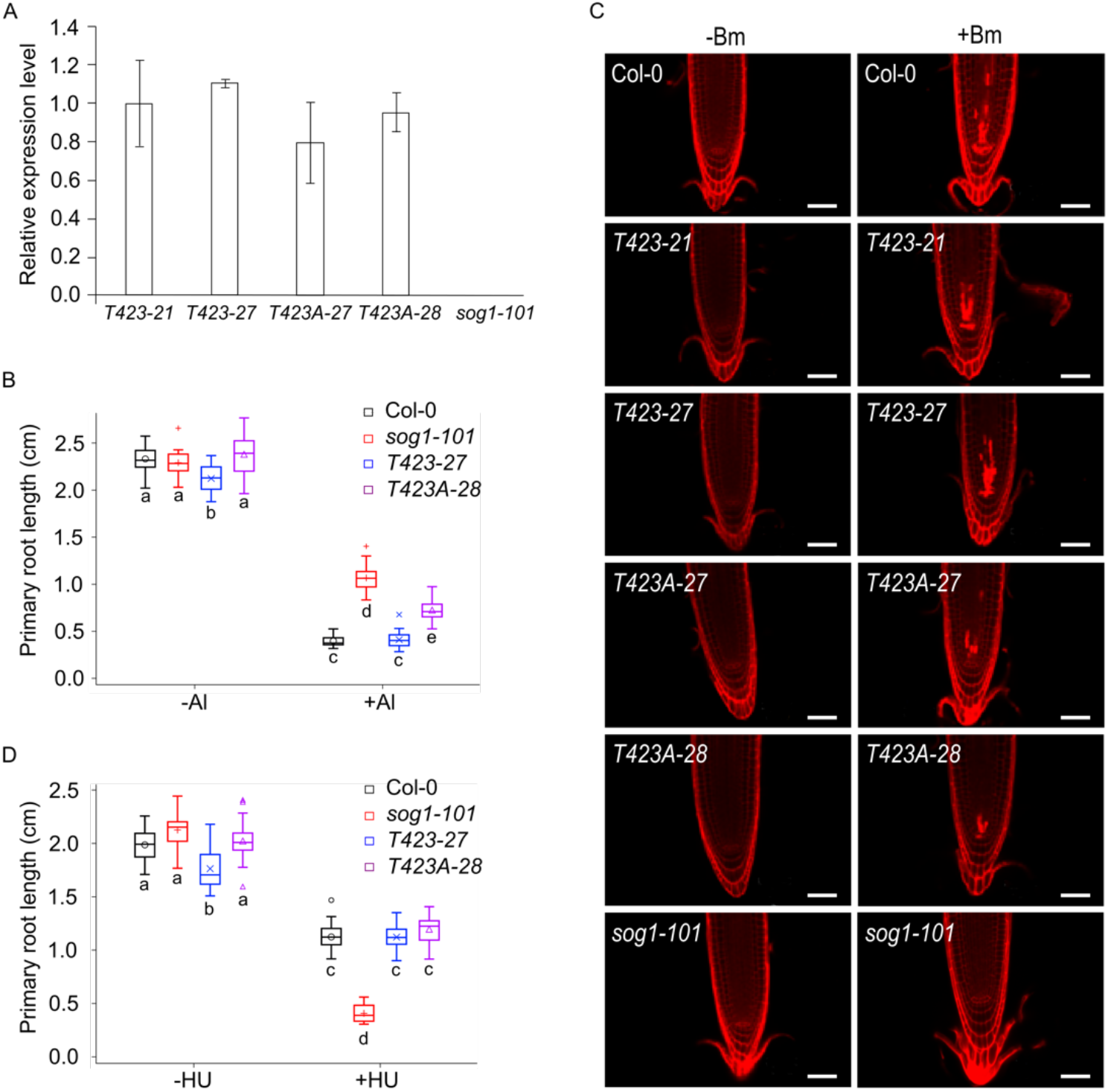
Molecular and Phenotypic Characterization *sog1-101* Complementation Lines (Supports Figure 7) (A) Relative *SOG1* expression levels in 5-day-old *sog1-101* seedlings and *sog1-101* seedlings holding a wild type (T423-21 and T423-27) or phospho-mutant (T423A-27 and T423A-28) *SOG1* allele. Data are represented by least square means ± se (n = 3). (B) Primary root length of wild-type (Col-0), *sog1-101*, T423-27 and T423A-28 plants grown for 7 days in a soaked gel environment (pH 4.2) in the absence (-Al) or presence of 1.5 mM AlCl_3_. (+ Al). Data are represented by box and whisker plots. Significant differences (p < 0.01) as determined by mixed model analysis with Tukey’s post-hoc correction are indicated by letters below bars (n = 3, with > 8 plants in each repeat). (C) Representative confocal microscopy images of meristems of 5-day-old seedlings transferred for 24 h to control medium (-Bm) medium or medium supplemented with 0.9 μg/ml bleomycin (+ Bm). Bars = 50μm. (D) Primary root length of wild-type, *sog1-101*, T423-27 and T423A-28 plants grown for 7 days on 1/2 MS control medium (-HU) or in the presence of 1 mM HU (+ HU). Data are represented by box and whisker plots. Significant differences (p < 0.01) as determined by mixed model analysis with Tukey’s post-hoc correction are indicated by letters below bars (n = 3, with > 8 plants in each repeat).

**Supplemental Figure 6.**
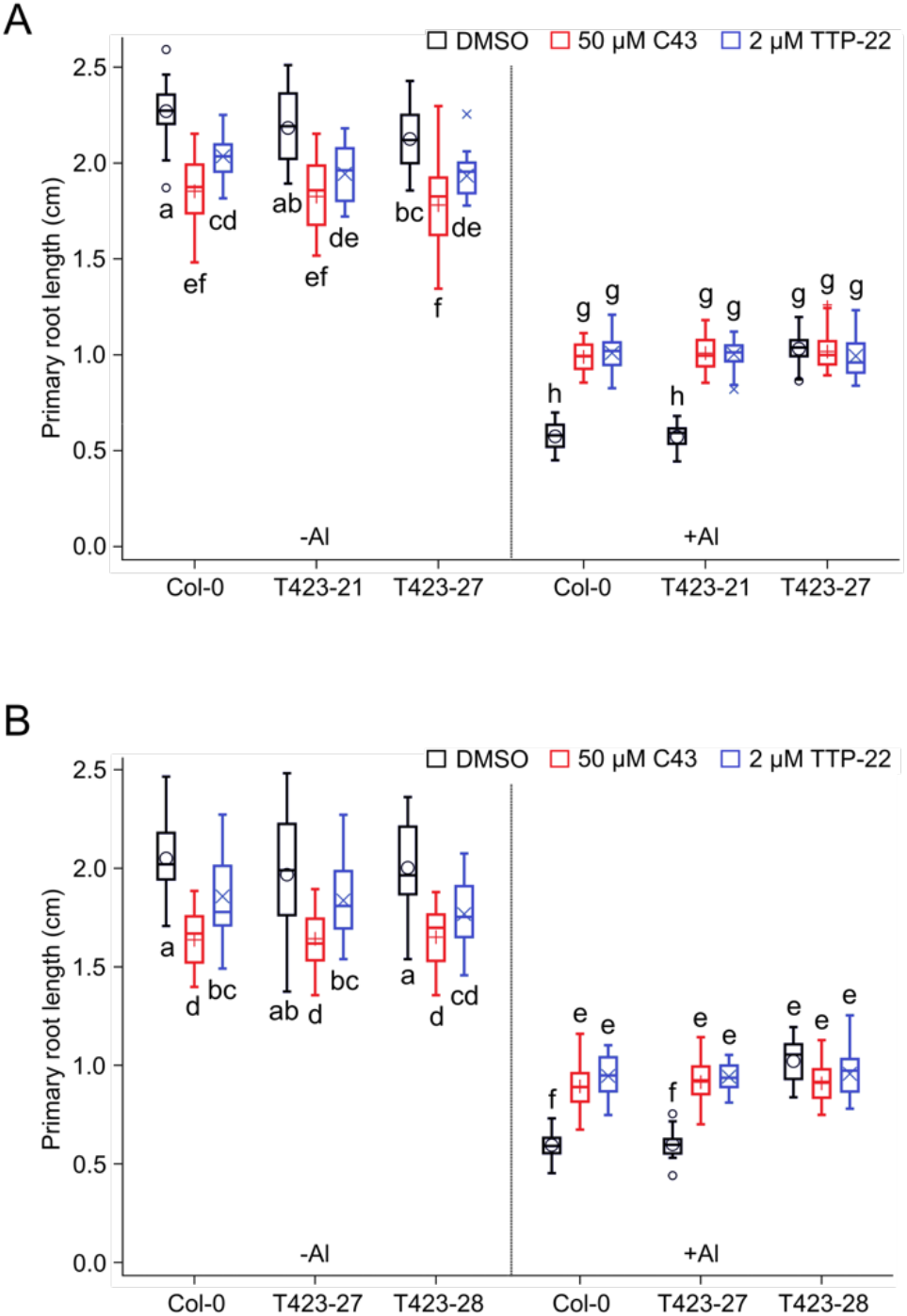
T423 represents a major CK2 phosphorylation site contributing to SOG1 activation (Supports Figure 7). Primary root length of 7-day-old wild-type (Col-0) and *sog1-101* complementation seedlings (T423-21 and T423A-27 (A) and T423-27 and T423A-28 (B)) grown in a soaked gel environment (pH 4.2) in the absence (-Al) or presence (+ Al) of 1.5 mM AlCl_3_, together with solvent control (DMSO), 50 μM C43 (A) or 2 μM TTP-22 (B). Data are represented by box and whisker plots. Significant differences (p < 0.05) as determined by mixed model analysis with Tukey’s post-hoc correction are indicated by letters above bars (n = 3, with > 8 plants in each repeat).

**Supplemental Figure 7.**
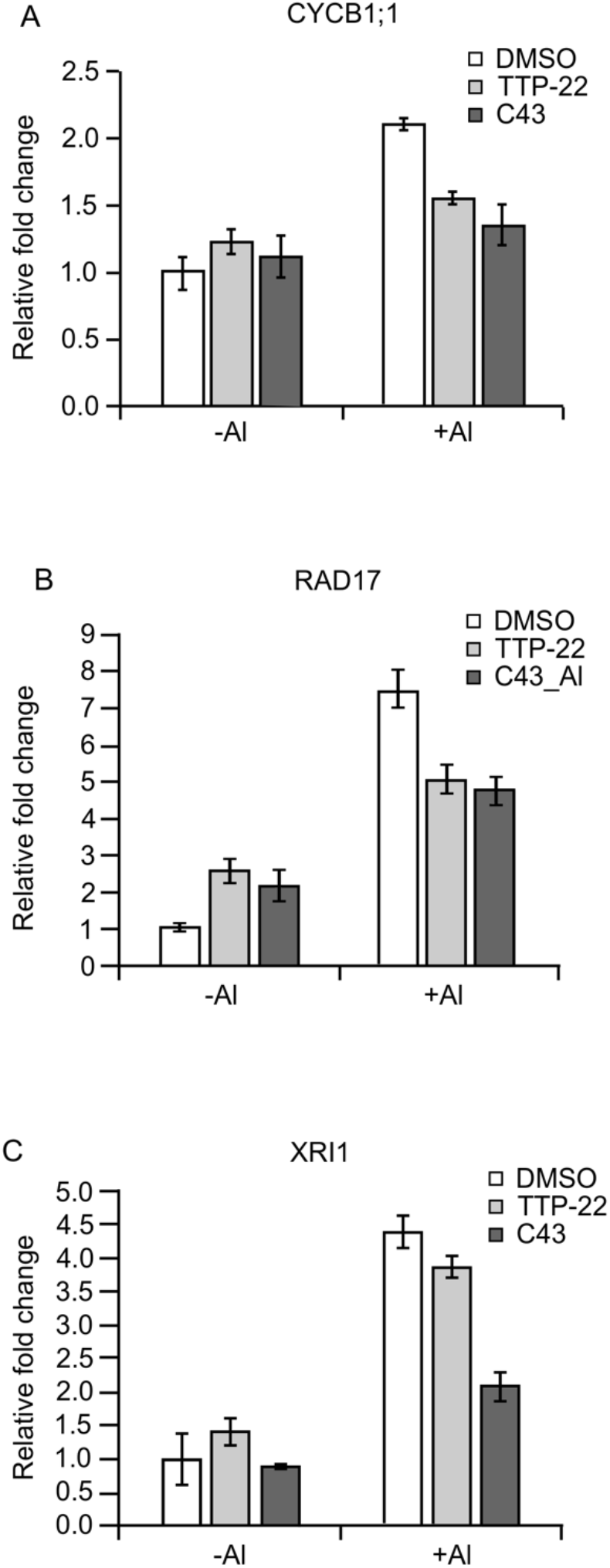
C43 and TTP-22 Repress Expression of SOG1 Target Genes (Supports Figure 7) Expression of *CYCB1;1* (A), *RAD17* (B) and *XRI1* (C) as measured by RT-qPCR in wild-type (Col-0) seedlings grown for 5 days (at pH 4.2) under control conditions (-Al) or in the presence of 1.50 mM AlCl_3_ (+ Al) together with solvent control (DMSO), 50 μM C43 or 2 μM TTP 22. The expression level of the DMSO control grown in the absence of Al was arbitrary set to one. Data represent least square means ± se (n = 3, with > 50 plants in each repeat).

**Supplemental Figure 8.**
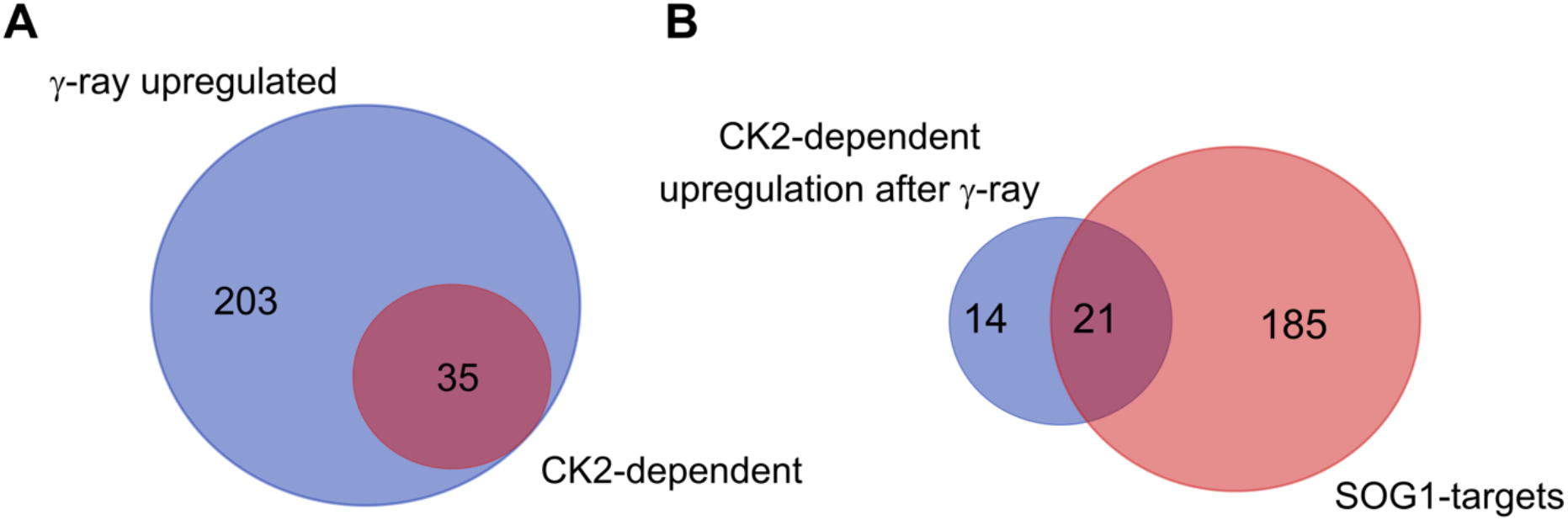
CK2-Dependent Upregulation of Genes Shows Statistically Significant Enrichment for SOG1 targets (Supports Figure 7) (A) Reanalysis of the transcriptome data published by Moreno-Romero et al. (2012), showing an upregulation of 238 genes upon γ-ray treatment, of which 35 are dependent on CK2. (B) Meta-analysis of the CK2-dependent genes upregulated after γ-ray treatment with a list of known SOG1 targets (Bourbousse et al., 2018) showing a statistically significant overlap (hypergeometric test, p < 0.001).

**Supplemental Figure 9.**
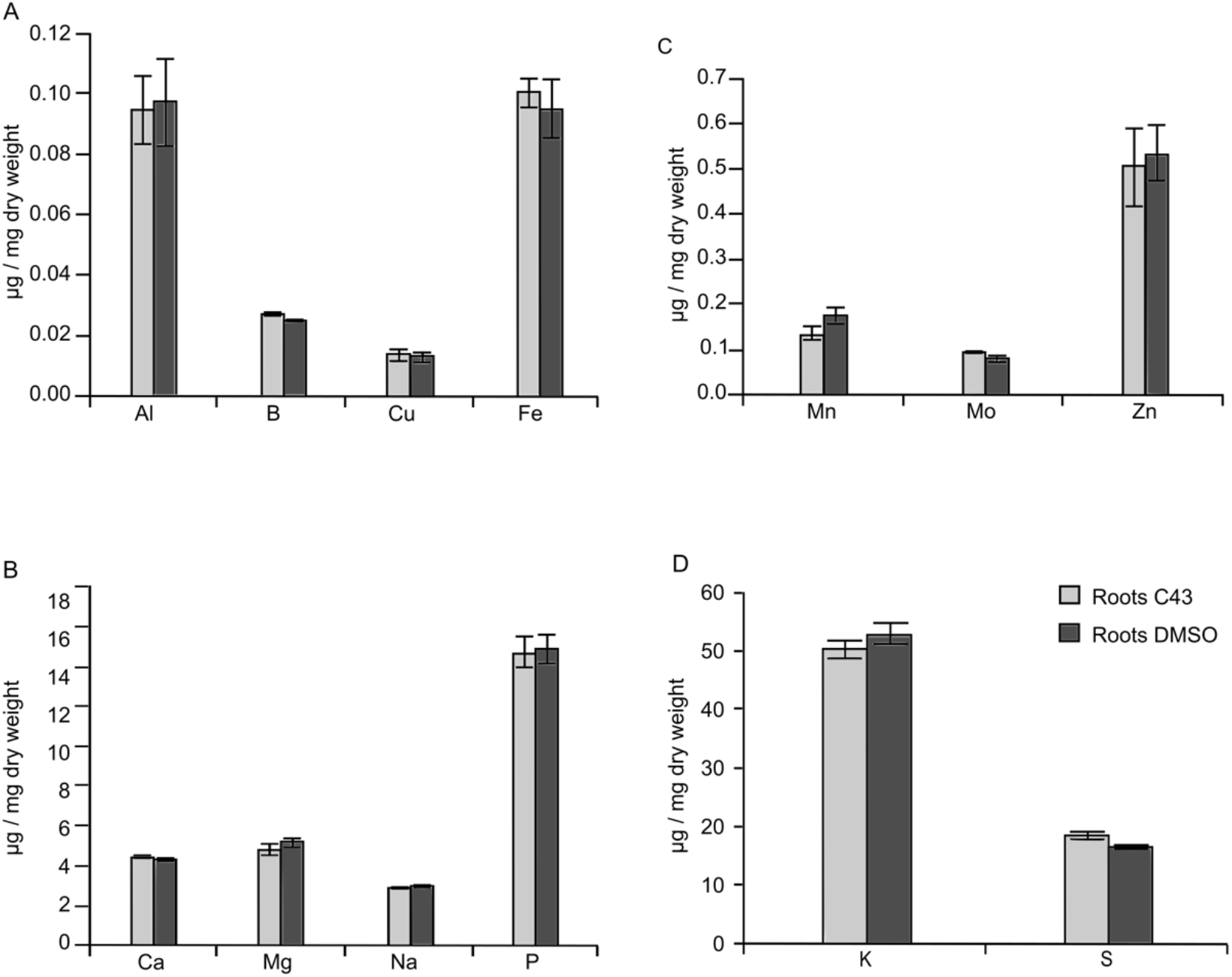
C43 Does Not Affect Mineral Composition of Roots (Supports Figure 9) ICP analysis of wild-type (Col-0) plants grown for 14 days in presence of the solvent control (DMSO) or 25 μM C43 for root 5 batches containing an average of 70 root system each. Data represent the average of two biological repeats.

